# Allosteric regulation of the tyrosine phosphatase PTP1B by a protein-protein interaction

**DOI:** 10.1101/2024.07.16.603632

**Authors:** Cassandra A. Chartier, Virgil A. Woods, Yunyao Xu, Anne E. van Vlimmeren, Andrew C. Johns, Marko Jovanovic, Ann E. McDermott, Daniel A. Keedy, Neel H. Shah

**Affiliations:** Department of Chemistry, Columbia University, New York, NY 10027; Structural Biology Initiative, CUNY Advanced Science Research Center, New York, NY 10031; Department of Biological Sciences, Columbia University, New York, NY 10027; PhD Program in Biochemistry, CUNY Graduate Center, New York, NY 10016; Department of Chemistry and Biochemistry, City College of New York, New York, NY 10031; PhD Programs in Biochemistry, Biology, & Chemistry, CUNY Graduate Center, New York, NY 10016

**Keywords:** Protein tyrosine phosphatase, PTP1B, allostery, Grb2, SH3 domain, protein-protein interaction, mass spectrometry

## Abstract

The rapid identification of protein-protein interactions has been significantly enabled by mass spectrometry (MS) proteomics-based methods, including affinity purification-MS, crosslinking-MS, and proximity-labeling proteomics. While these methods can reveal networks of interacting proteins, they cannot reveal how specific protein-protein interactions alter protein function or cell signaling. For instance, when two proteins interact, there can be emergent signaling processes driven purely by the individual activities of those proteins being co-localized. Alternatively, protein-protein interactions can allosterically regulate function, enhancing or suppressing activity in response to binding. In this work, we investigate the interaction between the tyrosine phosphatase PTP1B and the adaptor protein Grb2, which have been annotated as binding partners in a number of proteomics studies. This interaction has been postulated to co-localize PTP1B with its substrate IRS-1 by forming a ternary complex, thereby enhancing the dephosphorylation of IRS-1 to suppress insulin signaling. Here, we report that Grb2 binding to PTP1B also allosterically enhances PTP1B catalytic activity. We show that this interaction is dependent on the proline-rich region of PTP1B, which interacts with the C-terminal SH3 domain of Grb2. Using NMR spectroscopy and hydrogen-deuterium exchange mass spectrometry (HDX-MS) we show that Grb2 binding alters PTP1B structure and/or dynamics. Finally, we use MS proteomics to identify other interactors of the PTP1B proline-rich region that may also regulate PTP1B function similarly to Grb2. This work presents one of the first examples of a protein allosterically regulating the enzymatic activity of PTP1B and lays the foundation for discovering new mechanisms of PTP1B regulation in cell signaling.

**Significance Statement:** Protein-protein interactions are critical for cell signaling. The interaction between the phosphatase PTP1B and adaptor protein Grb2 co-localizes PTP1B with its substrates, thereby enhancing their dephosphorylation. We show that Grb2 binding also directly modulates PTP1B activity through an allosteric mechanism involving the proline-rich region of PTP1B. Our study reveals a novel mode of PTP1B regulation through a protein-protein interaction that is likely to be exploited by other cellular interactors of this important signaling enzyme.

## Introduction

Protein-protein interactions are central features of cell signaling pathways. They facilitate the colocalization of proteins within a cell, resulting in the coordination of protein function. In addition to increasing their local concentration, protein-protein interactions can also allosterically regulate protein function. Indeed, many signaling proteins operate via a switch-like mechanism, where the binding of one protein to another triggers its activation or inhibition.^1^ Recent advances in mass spectrometry proteomics have yielded a wealth of information about protein-proteins interactions that likely drive signaling.^2–8^ While these interactions are being catalogued with increasing speed and volume, the vast majority of them lack functional and structural characterization. Notably, it is unclear which of these protein-protein interactions mediate protein co-localization and which allosterically regulate protein function.

Protein tyrosine phosphatases are a family of enzymes that dephosphorylate tyrosine residues on proteins to modulate signaling processes. In humans, roughly 40 tyrosine phosphatases share a highly conserved catalytic domain, known as the classical protein tyrosine phosphatase domain, and many of these enzymes have additional non-catalytic domains that regulate their activity and localization. For example, the tyrosine phosphatase SHP2 has two Src homology 2 (SH2) domains, which control catalytic activity and localization in response to binding to tyrosine-phosphorylated proteins.^9^ The PDZ domain of the tyrosine phosphatase MEG1 recognizes protein C-termini in a sequence-specific manner, regulating MEG1 activity.^10,11^ By contrast, protein tyrosine phosphatase 1B (PTP1B), its paralog TCPTP, and other tyrosine phosphatases including BDP1 and LYP only have one globular domain: the catalytic domain (**Figure 1A**). Since these particular family members lack non-catalytic regulatory domains, it is less obvious how they might be regulated through protein-protein interactions, if at all. One possible mechanism is through interactions with their disordered C-termini, as in TCPTP.^12^ Alternatively, these phosphatases may be regulated through intermolecular interactions directly with the catalytic domain or flexible regions. Indeed, PTP1B is known to engage in a host of protein-protein interactions,^13–16^ not all of which involve substrates. Some of these interactors may allosterically modulate catalytic activity by binding distally to the active site and altering the structure or dynamics of the phosphatase.

**Figure 1.**
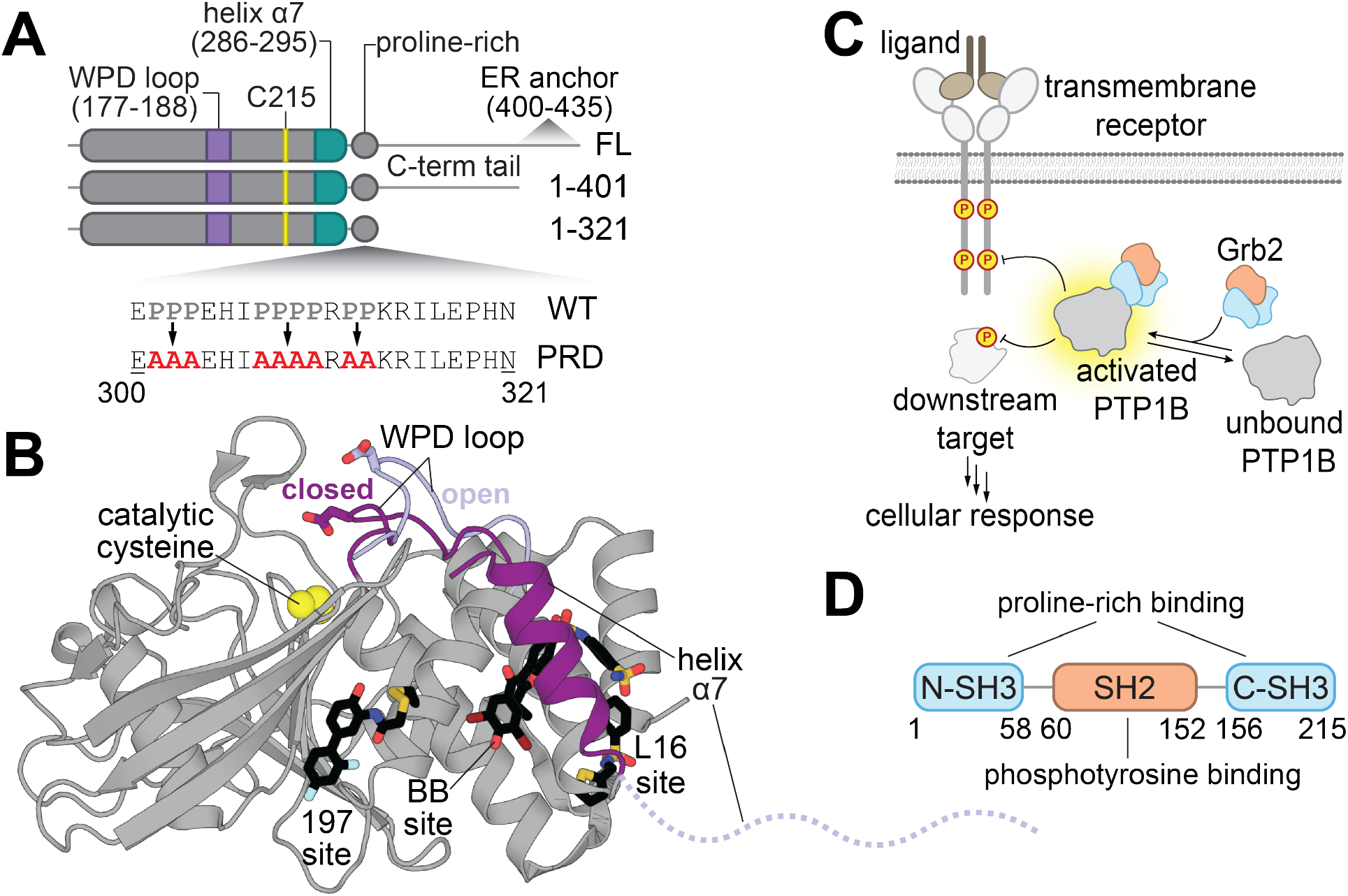
Structure and regulation of PTP1B. (**A**) Domain architectures and sequences of PTP1B constructs used in this study. Relevant structural features and mutagenized residues are indicated. (**B**) Overlay of active (PDB: 5K9W) versus inactive (PDB: 5K9V) PTP1B conformations. The active site is denoted by the catalytic residue Cys^215^ (yellow spheres). The active PTP1B conformation is characterized by a closed WPD loop and an ordered helix α7 (both dark purple). When PTP1B is inactive, the WPD loop opens (light purple) and helix α7 becomes disordered (dashed line). PTP1B can be allosterically inhibited by small molecules (black sticks) at the 197 site (PDB: 6B95), BB site (PDB: 1T49), and the L16 site (PDB: 5QFC). (**C**) The binding of Grb2 to PTP1B enhances its catalytic activity, which may impact relevant signaling pathways. (**D**) Domain architecture of Grb2.

Despite the dearth of evidence demonstrating that PTP1B can be allosterically regulated by protein-protein interactions, it is well-established that allostery is operative within the catalytic domain of this phosphatase.^17–20^ Studies of PTP1B allostery have been largely enabled by the discovery of small molecules that inhibit PTP1B activity by binding outside of the active site.^20–23^ These allosteric inhibitors bind at a few distinct sites, a subset of which are located near the conformationally dynamic helix α7 (**Figure 1B**). The conformation of α7 is coupled to the mobility of the catalytic WPD loop through a series of switchable non-covalent interactions between residues spanning almost 20 Å across the phosphatase domain. Given that small molecule binding can allosterically modulate PTP1B activity, we hypothesize that the engagement of endogenous protein interactors at allosteric sites might also alter PTP1B activity and regulate its function during cell signaling.

In this work, we examine whether a specific protein-protein interaction can allosterically regulate PTP1B activity (**Figure 1C**). We focus on the interaction of PTP1B with the adaptor protein Grb2, which has been previously studied in the context of insulin signaling.^24^ Grb2 consists of two terminal Src homology 3 (SH3) domains and a central SH2 domain (**Figure 1D**), which interact with proline-rich sequences and phosphotyrosine-containing sequences, respectively. As such, it has been suggested that the Grb2 SH3 domains can interact with the proline-rich region of PTP1B, which we confirm and quantitatively characterize in this work. Given that the proline-rich region is adjacent to helix α7 (**Figure 1A**), we hypothesized that binding to the proline-rich region might engage one of the known allosteric networks in PTP1B and alter catalytic activity.^17,20,25^ We show that PTP1B phosphatase activity is enhanced by Grb2 binding and provide evidence that this interaction alters PTP1B structure and/or dynamics using hydrogen-deuterium mass spectrometry (HDX-MS) and nuclear magnetic resonance (NMR) spectroscopy. We also show that Grb2 binding to PTP1B via the proline-rich region can occur in cells, and we use mass spectrometry proteomics to identify other proteins that bind at this site, yielding an array of potential regulators of PTP1B activity. We envision that this work will set the stage for the characterization of other PTP1B interactors and serve as baseline evidence for the allosteric regulation of classical tyrosine phosphatase domains through interactions with endogenous proteins.

## Results

### The proline-rich region of PTP1B is essential for Grb2 binding

Previous work has shown that the proline-rich region of PTP1B is the primary binding site for the SH3 domain-containing protein p130cas.^13^ The isolated Grb2 SH3 domains were also previously shown to bind PTP1B, but the binding site on PTP1B was not determined.^13^ We hypothesized that Grb2 also binds to the proline-rich region of PTP1B via its SH3 domains. To test this hypothesis, we carried out an array of binding experiments using various PTP1B and Grb2 constructs. For most experiments, we used the Grb2^Y160E^ mutant because of its high solubility and low propensity for dimerization relative to Grb2^WT^,^26,27^ but key findings that we obtained using Grb2^Y160E^ were corroborated using Grb2^WT^. First, we used size-exclusion chromatography to assess the binding of Grb2 to PTP1B (**Figure 2A**). We observed a shift in elution volume for a mixture of wild-type PTP1B^1-321^ and Grb2^Y160E^, indicating complex formation. This was further confirmed through SDS-PAGE analysis of the elution fractions, which showed co-elution of Grb2^Y160E^ with PTP1B^1-321^ (**Figure 2A**). Based on the co-elution volume of PTP1B and Grb2 in the mixture, the PTP1B-Grb2 complex most likely has a 1:1 stoichiometry under these experimental conditions (**Figure S1A**). We cannot rule out the possibility that PTP1B and Grb2 form higher-order complexes at higher concentrations.

**Figure 2.**
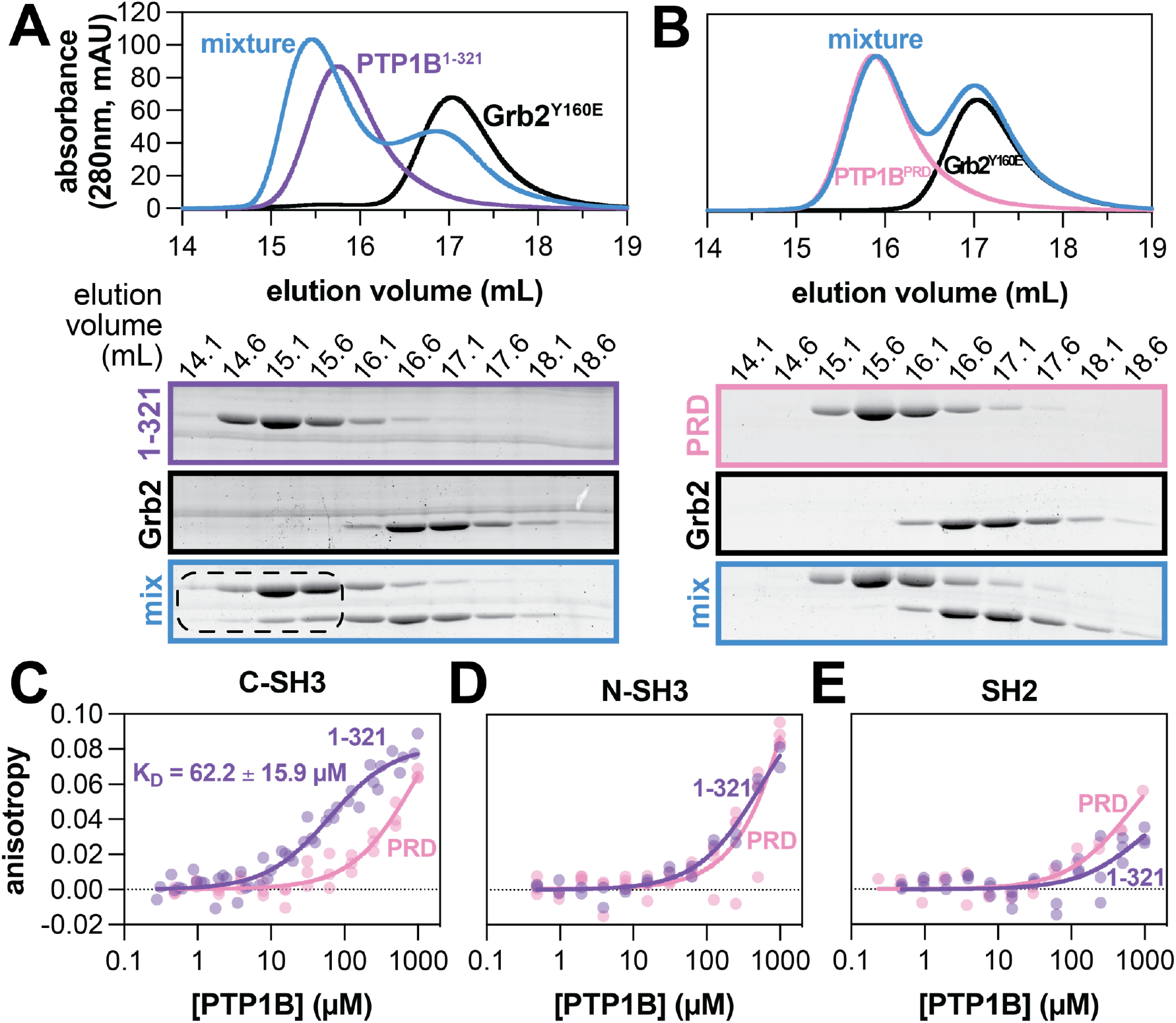
Grb2 binding to PTP1B is mediated by interaction of the PTP1B proline-rich region and Grb2 C-SH3 domain. (**A**) Size exclusion chromatograms (top) depicting separate injections of equimolar PTP1B^1-321^, Grb2^Y160E^, and a mixture of the two proteins (20 μM each). SDS-PAGE analysis (bottom) shows co-elution of PTP1B^1-321^ and Grb2^Y160E^ at earlier elution volumes in the mixture. (**B**) Size exclusion chromatograms (top) depicting separate injections of equimolar PTP1B^PRD^, Grb2^Y160E^, and a mixture of the two proteins (20 μM each). SDS-PAGE analysis (bottom) shows the proteins in the mixture elute at the same volumes as when not in a mixture. (**C-E**) Fluorescence anisotropy analysis of PTP1B binding to fluorescently labeled isolated Grb2 domains. N = 2-4 independent titrations of PTP1B with fluorescently labeled isolated Grb2 domain. Data points from all replicates are plotted on each graph.

Next, we assessed the importance of the proline-rich region for binding by mutating the proline residues to alanine (**Figure 1A**). This proline-rich dead construct (PTP1B^PRD^) did not co-elute with Grb2^Y160E^ at an earlier elution volume than the individual proteins, suggesting a lack of complex formation (**Figure 2B**). These trends were also observed using Grb2^WT^, indicating that complex formation is not dependent on the Y160E mutation (**Figure S1B,C**). We also tested whether the disordered tail of PTP1B impacted its ability to bind Grb2, as this tail has previously been shown to regulate PTP1B activity.^28^ Size-exclusion chromatography analysis of Grb2^WT^ mixed with PTP1B^1-401^, which includes the disordered tail, confirmed that these two proteins can form a complex (**Figure S1D**). Overall, these experiments confirm that the proline-rich region is essential for the PTP1B-Grb2 interaction and that the disordered tail of PTP1B does not impede this interaction.

We then sought to quantify the affinity of the PTP1B-Grb2 interaction and determine which Grb2 domain drives binding to PTP1B, using fluorescence anisotropy measurements. First, we labeled the N-termini of full-length Grb2 and each isolated domain with a fluorophore, using a sortase-mediated ligation reaction (**Figure S2A**).^29^ Then, we titrated PTP1B into a solution of each labeled Grb2 construct and measured fluorescence anisotropy. Given the similar size of full-length Grb2 and PTP1B^1-321^, we were unable to obtain high-quality anisotropy data for this construct, but we estimate the dissociation constant (*K*_D_) to be in the 20-30 μM range (**Figure S2B**). By contrast, the smaller isolated domains yielded high-quality fluorescence anisotropy data (**Figure 2C-E**). PTP1B^1-321^ exhibited the strongest binding to the Grb2 C-SH3 domain with a dissociation constant of 62 ± 16 μM, which was severely attenuated by mutating the proline-rich region (**Figure 2C**). Robust binding to the C-SH3 domain was also observed with PTP1B^1-401^, with a comparable affinity of 82.4 ± 4.3 μM (**Figure S2C**). Interestingly, titration of PTP1B with the Grb2 N-SH3 domain demonstrated a measurable increase in anisotropy at high PTP1B concentrations, independent of the proline-rich region (**Figure 2D**), suggesting a possible weak secondary binding site. The C-SH3 domain showed a similar signal at high PTP1B concentrations, suggesting that it may also weakly bind PTP1B at a site other than the proline-rich region (**Figure 2C**). PTP1B titration with the Grb2 SH2 domain exhibited the weakest change in anisotropy of the individual domains (**Figure 2E**), indicating minimal or no role in mediating Grb2 binding to PTP1B. Taken together, these results show that the C-SH3 domain is the key player in Grb2 binding to PTP1B. Furthermore, they corroborate the importance of the proline-rich region for Grb2 binding to PTP1B.

#### Grb2 binding alters PTP1B activity against phosphopeptide substrates

To understand the functional consequences of Grb2 binding to PTP1B, we examined its effect on PTP1B catalytic activity. Previous work has shown that PTP1B-mediated dephosphorylation of the protein IRS-1, which is tyrosine-phosphorylated at many sites, is enhanced in the presence of Grb2.^24^ In that prior study, enhanced dephosphorylation was attributed to increased co-localization and not to allosteric activation of PTP1B. This hypothesis was based on the idea that Grb2 could bridge PTP1B and IRS-1 via interactions between the SH3 domains and PTP1B as well as the SH2 domain and phosphosites on IRS-1. To exclude contributions from co-localization and directly assess if Grb2 allosterically activates PTP1B, we measured the dephosphorylation kinetics of mono-phosphorylated peptide substrates by PTP1B in the presence and absence of Grb2^Y160E^ (**Figure 3A and Figure S3A**). We selected a series of peptide sequences derived from proteins that are known substrates of PTP1B or involved in PTP1B-related pathways, including Gab1, EGFR, c-Src, and IRS-1.^24,30–34^ For every peptide tested, the presence of Grb2^Y160E^ enhanced *k*_cat_ (**Figure 3B**), suggesting that Grb2 binding to PTP1B stabilizes a catalytically-competent conformation of the phosphatase. Grb2^Y160E^ alone does not have any measurable ability to dephosphorylate substrates in this experiment, as expected (**Figure S3B**). Surprisingly, Grb2 binding to PTP1B caused varying effects on *K*_M_ (**Figure S3C**), indicative of changes in substrate recognition. In a cellular context, where signaling proteins are often co-localized with their substrates through auxiliary protein-protein interactions, the effect of *k*_cat_ likely dominates over *K*_M_. While the overall effects on catalytic activity were modest, we note that previous studies investigating altered PTP1B activity have shown that small changes in activity are sufficient to elicit large changes in phosphorylation of downstream proteins.^35–37^

**Figure 3.**
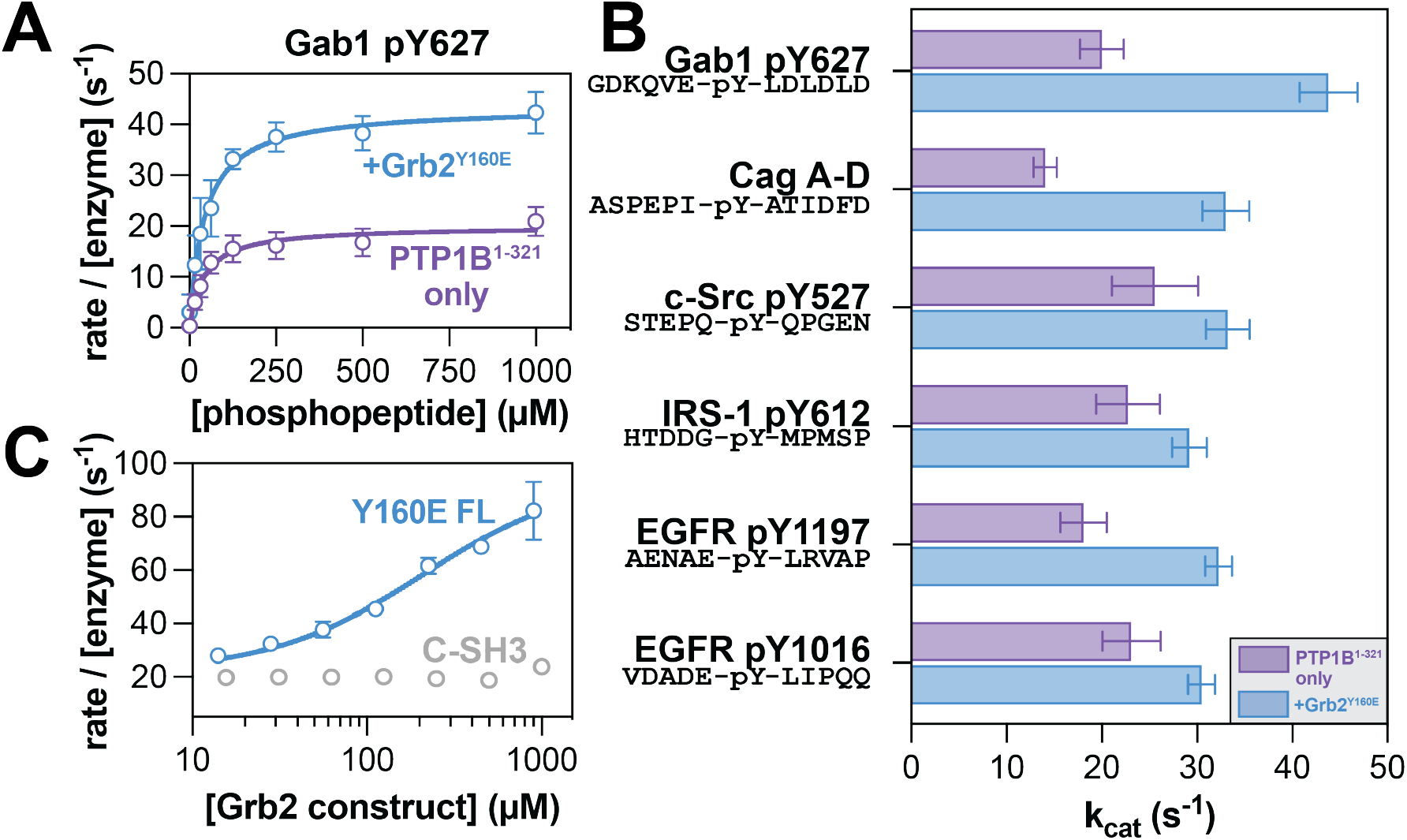
Grb2 enhancement of PTP1B activity. (**A**) Representative Michaelis-Menten curve showing PTP1B^1-321^ activity alone (purple) and in response to 50 µM Grb2 Y160E (blue) against the Gab1 phosphopeptide substrate. N = 3-4 independent titrations of Gab1. (**B**) Measured *k*cat values for PTP1B against all phosphopeptide substrates tested. (**C**) PTP1B activity against Gab1 phosphopeptide substrate in response to increasing concentrations of Grb2^Y160E^ (blue) and Grb2 C-SH3 domain (grey). N = 2-3 independent titrations of full-length Grb2^Y160E^ or isolated C-SH3 domain.

Next, we examined the effects of Grb2-mediated PTP1B activation with various PTP1B and Grb2 constructs. First, we confirmed that full-length Grb2^WT^ could activate PTP1B^1-321^, demonstrating that this effect is not specific to the Y160E mutation (**Figure S3D**). We further confirmed that PTP1B^1-401^ is also activated by Grb2^WT^, indicating that the disordered tail of PTP1B does not impact this allosteric effect (**Figure S3E**). We also assessed whether the isolated C-SH3 domain of Grb2 was sufficient for activation, given that its binding to the PTP1B proline-rich region drives the interaction of these two proteins. PTP1B^1-321^ activity against the Gab1 phosphopeptide showed a titratable increase with the addition of full-length Grb2^Y160E^, with an approximate EC_50_ value of 215 μM (**Figure 3C**). This is consistent with our measured mid-micromolar *K*_D_ for the Grb2 C-SH3 domain and PTP1B^1-321^ (**Figure 2C**). By contrast, titration of the isolated C-SH3 domain of Grb2 did not enhance of PTP1B^1-321^ activity (**Figure 3C**). This suggests that the C-SH3 domain is necessary for binding, but not sufficient for the allosteric effect on PTP1B activity. Our fluorescence anisotropy data suggest that the N-SH3 domain might weakly bind to PTP1B at a site distinct from the proline-rich region (**Figure 2D**). Thus, one plausible explanation for our functional data is that the C-SH3 domain acts as a tether, allowing the N-SH3 domain to contact PTP1B and potentiate the allosteric effect on activity observed with full-length Grb2. Alternatively, we note that previous work on Grb2 has demonstrated that the individual domains can allosterically modulate one another’s binding functions, which could explain why the isolated C-SH3 domain can bind to but not activate PTP1B.^38,39^

#### Grb2 binding alters PTP1B structure and dynamics

Given that Grb2 binding to PTP1B enhances catalytic activity, we explored whether this was caused by a change in PTP1B structural dynamics. To probe this, we first used global HDX-MS to assess whether Grb2 binding to PTP1B altered the deuterium uptake of PTP1B, which would suggest a change in its dynamics (**Figure 4A**). Upon titrating Grb2^Y160E^ into a sample of PTP1B, we observed a decrease in deuteration, consistent with an interaction between the two proteins (**Figure 4B and Figure S4A**). This result is consistent with blocking H/D exchange of non-proline residues in and around the proline-rich region, which is the primary Grb2 binding site. It is also possible that Grb2 binding reduces dynamics elsewhere in PTP1B and/or Grb2 makes contact with another site on the PTP1B surface. Notably, the effect of Grb2^Y160E^ on PTP1B H/D exchange was mostly recapitulated using the isolated C-SH3 domain (**Figure 4B**). Furthermore, we did not observe a significant effect on H/D exchange using the isolated N-SH3 domain (**Figure 4B**), nor was there an effect of Grb2^Y160E^ binding on H/D exchange in PTP1B^PRD^ (**Figure S4B**). These results are consistent with our binding and biochemical measurements, which show that the C-SH3 domain is the primary contributor to PTP1B binding.

**Figure 4.**
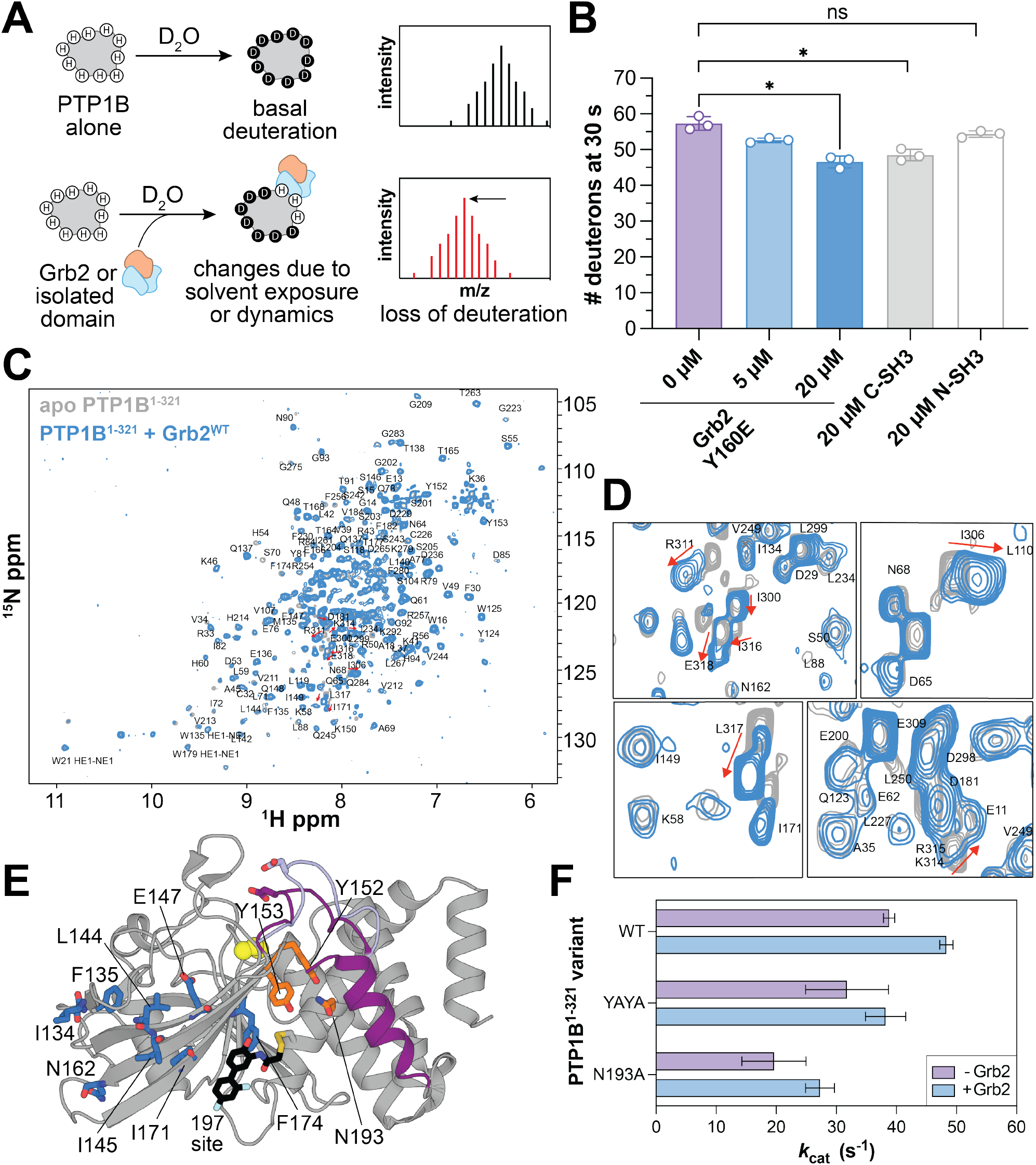
Changes to structure and/or dynamics induced by Grb2 binding to PTP1B. (**A**) Schematic of global HDX-MS experiments. (**B**) Deuterium exchange of PTP1B^1-321^ as Grb2^Y160E^ is added. A paired, two-tailed t-test was used to test for significance (* denotes p < 0.05, ns denotes p > 0.05 or not significant). (**C**) Overlay of apo PTP1B^1-^ 321 (grey) and Grb2^WT^-bound PTP1B^1-321^ (blue) ^1^H-^15^N HSQC spectra. Assigned peaks are labeled by residue. (**D**) Chemical shift changes within the proline-rich region induced by Grb2 binding. Peaks corresponding to K314, R311, I300, I316, E318, I306, and L317 (top to bottom) are indicated. See Figure 1B for proline-rich sequence. (**E**) Residues with minor chemical shift or peak changes are located in a β-sheet near the proline-rich region (blue sticks). Some residues involved in allosteric communication with helix α7 (Y152, Y153, and N193) are indicated. (**F**) *k*cat for PTP1B^1-321^ variants (wild-type, YAYA, and N193A) with the Gab1 phosphopeptide substrate, with and without Grb2^Y160E^. N = 3 independent titrations of Gab1.

To gain additional structural insights into the PTP1B-Grb2 interaction, we used NMR spectroscopy. We obtained ^1^H-^15^N HSQC spectra of ^15^N-labeled PTP1B^1-321^ alone and in complex with unlabeled Grb2^WT^. The spectrum of isolated PTP1B^1-321^ aligns closely with a previously published dataset of apo PTP1B (BMRB ID: 19224),^28^ enabling confident assignment of many backbone amide N-H peaks (**Figure S4C and Table S1**). However, there were significant peak shifts or disappearances in our spectrum corresponding to residues near the active site, such as such as G86, L110, R112, V113, C215, G218, I219, G259, and L260. These residues are known to be flexible in the absence of a substrate or inhibitor and were detectable in the published dataset due to deuteration and TROSY-HSQC detection, which reduces spin relaxation.

When comparing the spectra of PTP1B^1-321^ alone and in complex with Grb2^WT^ (**Figure 4C**), we observed pronounced chemical shift changes for non-proline residues within and immediately adjacent to the proline-rich region of PTP1B, namely I300, I306, R311, K314, R315, I316, L317, and E318 (**Figure 1A and Figure 4D**). This further supports the notion that the proline-rich region is the primary Grb2 binding site, as demonstrated by our binding studies (**Figure 2A-C**). The perturbations at R311 and K314 are particularly noteworthy, as the Grb2 C-SH3 domain has been shown to have non-canonical specificity by recognizing RXXK motifs, where X can be any amino acid, including proline.^40^ Consistent with this, a previous study examining a dozen different SH3 domains showed that most of these do not bind PTP1B, suggesting some specificity to this interaction.^13^ Additionally, we noted minor chemical shift perturbations or peak intensity changes for residues in the main β-sheet that runs through the phosphatase domain (**Figure S4D**). Interestingly, a subset of these residues, namely L144, I145, E147, I171, and F174, are located near a recently discovered allosteric covalent inhibitor binding site known as the “197 site” (**Figure 4E**).^20^

Finally, we examined whether the effects of Grb2 binding on PTP1B activity are exerted through one of the known allosteric mechanisms that is exploited by small molecule inhibitors. In particular, given the proximity of helix α7 to the proline-rich region, we wondered if helix α7 is central to this effect. In previous reports, mutations near helix α7 have been designed that disrupt the interaction between the globular domain and ordered helix α7, thereby weakening allosteric communication.^17^ We evaluated whether Grb2 could still exert its allosteric effect on PTP1B in the context of two of these mutants: one in which Y152 and Y153, are simultaneously mutated to alanine (PTP1B^YAYA^), and another where N193 is mutated to alanine (PTP1B^N193A^) (**Figure 4E**). In past work, both mutations have been shown to slightly decrease basal PTP1B activity toward the small molecule substrate *p*-nitrophenyl phosphate.^17^ A similar decrease in activity was observed in our phosphopeptide dephosphorylation assays. Notably, the activity-enhancing effect of Grb2^Y160E^ was still observed with PTP1B^YAYA^ and PTP1B^N193A^ (**Figure 4F and Figure S5**). These results suggest that Grb2 activates PTP1B through a distinct allosteric mechanism that is not driven by interactions between helix α7 and the globular phosphatase domain.

#### The proline-rich region of PTP1B is a binding hotspot for other proteins in cells

We have shown that the proline-rich region of PTP1B is a regulatory handle for Grb2 in a biochemical context. To demonstrate the importance of this region for the PTP1B-Grb2 interaction in cells, we transfected HEK 293 cells with plasmids encoding full-length PTP1B^WT^ or PTP1B^PRD^ containing an N-terminal myc-tag. PTP1B was immunoprecipitated using myc-tag-specific beads, and levels of co-immunoprecipitated endogenous Grb2 were evaluated by western blot (**Figure 5A**). As expected, Grb2 co-immunoprecipitated with PTP1B^WT^, but not with PTP1B^PRD^ (**Figure 5B**). We repeated this experiment for another SH3 domain-containing protein, ARHGAP12, that has been identified in proteomics studies as a PTP1B interactor, but for which the binding site on PTP1B has not been determined.^14^ Co-immunoprecipitation of this interactor was also dependent on the proline-rich region (**Figure 5C**). These results show that the proline-rich region is important for both Grb2 and ARHGAP12 binding to PTP1B in cells. Furthermore, they demonstrate that the PTP1B-Grb2 interaction remains intact in the context of full-length PTP1B. Notably, the PTP1B construct used in these experiments contains the C-terminal tail and ER membrane anchor (**Figure 1A**), which allows PTP1B to at least partially retain its ER localization.^41,42^

**Figure 5.**
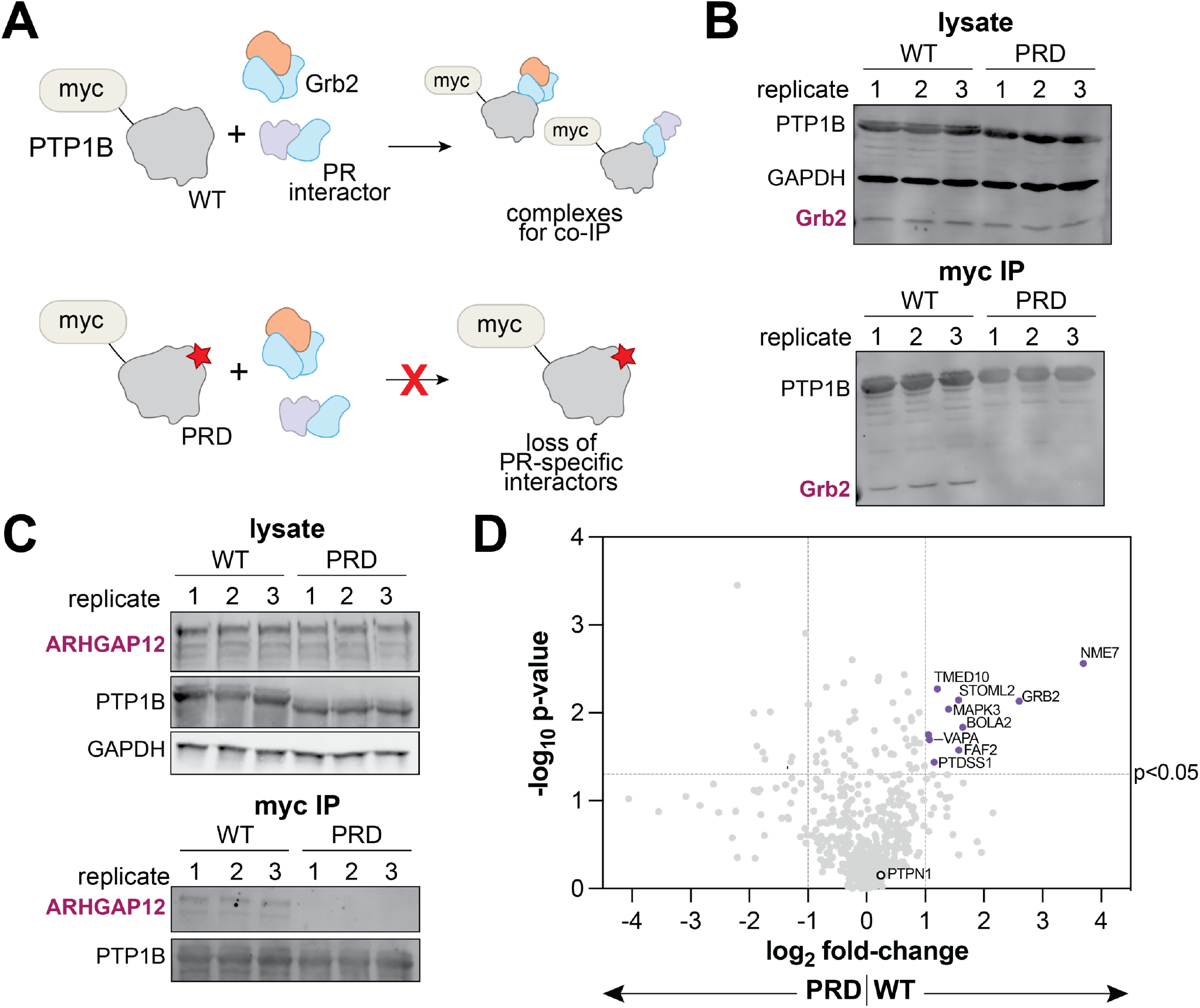
PTP1B proline-rich region interactors identified by proteomics. (**A**) Schematic showing the co-immunoprecipitation of interacting proteins with PTP1B. PTP1B^WT^ interacts with Grb2 and other proteins specific for the proline-rich region, while PTP1B^PRD^ will not pull down the same interactors due to its mutagenized proline-rich region. (**B**) Grb2 co-immunoprecipitation with PTP1B^WT^ and PTP1B^PRD^. Replicates are from six separate transfections are analyzed on the same western blot. (**C**) ARHGAP12 co-immunoprecipitation with PTP1B^WT^ and PTP1B^PRD^. Replicates are from six separate transfections are analyzed on the same western blot. (**D**) Volcano plot highlighting putative interactors of the PTP1B proline-rich region (purple dots).

Given the importance of the proline-rich region for the PTP1B-Grb2 interaction in cells, we hypothesized that this region may also be implicated in other regulatory protein-protein interactions. Thus, we immunoprecipitated myc-tagged PTP1B^WT^ or PTP1B^PRD^ from HEK 293 cell lysates and identified the enriched proteins by mass spectrometry proteomics. As expected, PTP1B, encoded by the *PTPN1* gene, was detected with the highest intensity in all samples, confirming successful immunoprecipitation via the myc-tag (**Figure S6**). Many proteins were selectively enriched in the PTP1B^WT^ samples over the PTP1B^PRD^ samples, suggesting that those proteins engage PTP1B via the proline-rich region (**Figure 5D and Table S2**). Critically, Grb2 was enriched in all PTP1B^WT^ replicates but undetectable in all PTP1B^PRD^ replicates (**Figure 5D and Figure S6**), in agreement with our co-immunoprecipitation experiments (**Figure 5B**). We were surprised to find that none of the hits, other than Grb2, contained an SH3 domain or other canonical polyproline binding domain, suggesting alternative modes of recognition that warrant further structural studies. Notably, the SH3-containing protein ARHGAP12, which we showed selectively co-immunoprecipitated with PTP1B^WT^ over PTP1B^PRD^ (**Figure 5C**), was present in our PTP1B^WT^ proteomics samples and not detected in the corresponding PTP1B^PRD^ samples. However the abundance of this protein was too low in our proteomics datasets to be confidently deemed a hit.

Interestingly, we found that several proteins enriched by PTP1B^WT^ over PTP1B^PRD^ are localized to the ER (FAF2, PTDSS1, TMED10, and VAPA), suggesting that the proline-rich region may be a binding site for proteins co-localized with PTP1B. One of these proteins, VAPA, was previously found to interact with PTP1B in a proximity-labeling proteomics study.^43^ Both VAPA and PTP1B play a role in the formation of focal adhesions and promote cell migration.^44,45^ While VAPA does not have a canonical polyproline binding domain, it is known to bind to sequences containing diphenylalanine in an acidic tract (FFAT) motifs.^46^ A few residues after the proline-rich region, PTP1B has a diphenylalanine motif preceded by a glutamate residue, which could be a binding site for VAPA that is disrupted in PTP1B^PRD^. Overall, the structural hypothesis-driven proteomics experiments described in this section demonstrate that many proteins likely interact with PTP1B via its proline-rich region. It is plausible that some of these proteins may regulate activity through a mechanism similar to that of Grb2.

## Discussion

Several recent proteomics studies have revealed a suite of protein interactors for the tyrosine phosphatase PTP1B.^14,15,47^ The functional consequences of many of these non-substrate PTP1B interactions remain unknown. Indeed, very few proteins have been shown to directly regulate PTP1B activity.^48^ In this study, we investigated the interaction between PTP1B and the adaptor protein Grb2. Previously, these proteins were suggested to form a ternary complex with the PTP1B substrate IRS-1, leading to enhanced dephosphorylation through co-localization.^24^ In this work, we show that Grb2 binding to PTP1B directly enhances phosphatase activity, independent of auxiliary interactions that co-localize the substrate with PTP1B. We pinpoint this interaction to the proline-rich region of PTP1B and the C-SH3 domain of Grb2, and we show that the interaction between PTP1B and Grb2 can be recapitulated in mammalian cell culture.

In our biophysical studies, we observed that Grb2 binding to PTP1B structurally perturbs the phosphatase. However, our results leave a few open questions about the allosteric mechanism. We note that the isolated C-SH3 domain is competent to bind to PTP1B, but not sufficient to allosterically alter its catalytic activity. In our H/D exchange experiments, there is a modest difference between full-length Grb2 and the isolated C-SH3 domain, suggesting there are additional structural perturbations occurring upon binding of the full-length construct. It is important to note that the linker between the C-SH3 and SH2 domain, which is not present in the isolated C-SH3 domain construct, could play a role in the allosteric effect. Another possible explanation is that the N-SH3 domain forms a second weak interaction with PTP1B that results in the allosteric effect caused by binding of full-length Grb2. Indeed, our binding studies revealed a weak interaction between the N-SH3 domain and PTP1B that is independent of the proline-rich region.

There are no high-resolution structures of Grb2 bound to PTP1B. Thus, we generated models using AlphaFold 3 to guide our speculation about the modes of interaction and mechanisms of allostery.^49^ Most of the models showed either the N-or C-SH3 domain engaging the proline-rich region of PTP1B. Based on our binding studies, we ruled out models where the N-SH3 domain was bound to the proline-rich region. We also ruled out models where the active site was occluded, which would be inconsistent with our activity studies. The remaining model shows the N-SH3 domain forming an interface above the WPD loop, in a region where it may alter catalytic activity (**Figure 6A**). Although this N-SH3 interface does not resemble a canonical SH3-peptide interaction, it is noteworthy that the same region in the SHP2 phosphatase domain was recently suggested to bind to the Grb2 C-SH3 domain.^26^ In our AlphaFold 3 model, the C-SH3 domain not only binds the proline-rich region, but it also forms an interface with the E loop. This loop has correlated motions with the WPD loop, and it is also important for substrate recgnition, which may explain why Grb2 has varied effects on PTP1B depending on the substrate sequence (**Figure 3B and Figure S3C**).^18,50,51^ Given these observations and our biochemical data, we propose two plausible mechanisms for the Grb2-mediated enhancement of PTP1B activity in which (i) the N- and C-SH3 domains bind to distinct sites on PTP1B or (ii) only the C-SH3 domain makes direct contact with PTP1B, but its binding is modulated by interdomain allostery within Grb2 (**Figure 6B**).

**Figure 6.**
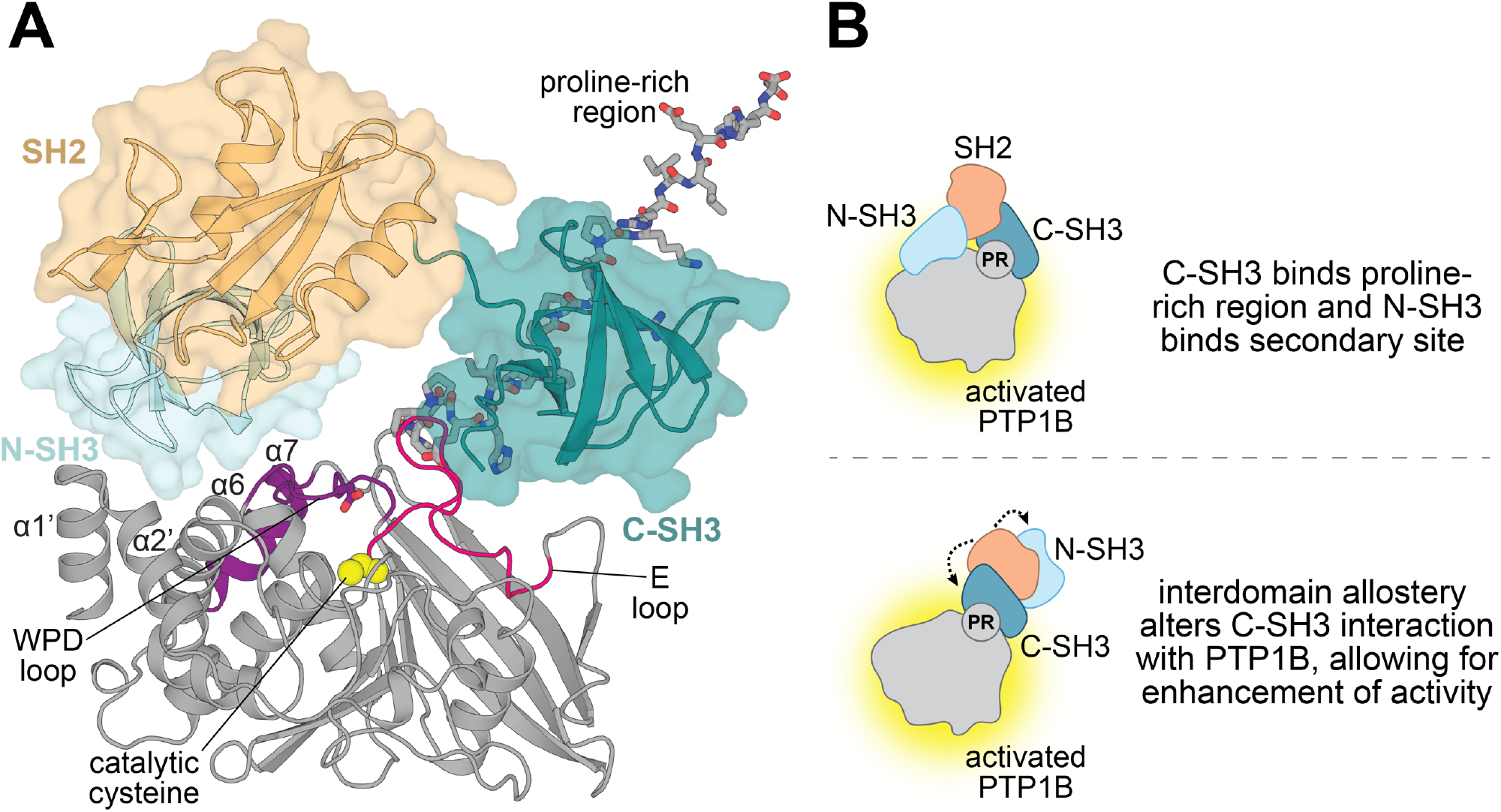
PTP1B-Grb2 complex structural model and proposed allosteric mechanism. (**A**) AlphaFold 3-generated model of PTP1B^1-321^ (grey) bound to full-length Grb2^WT^. The proline-rich region is shown as grey sticks. The Grb2 C-SH3 domain (deep teal) is shown interacting with the proline-rich region and the E loop (hot pink). The N-SH3 domain (light cyan) is shown forming an interface with the N-terminal helices α1’ and α2’, helix α6, and the WPD loop. The WPD loop is in the closed conformation in this model. The SH2 domain is pointed away and does not form any interfaces with PTP1B. (**B**) Proposed mechanisms for allosteric enhancement of PTP1B activity by Grb2. Grb2 C-SH3 domain binds the proline-rich region while the N-SH3 domain interacts with a secondary site, allowing Grb2 to allosterically enhance PTP1B activity (top). Grb2 C-SH3 domain allosterically enhances PTP1B activity as a result of interdomain allostery in Grb2, which alters the interaction between the C-SH3 domain and PTP1B (bottom).

Much of what we know about PTP1B allostery is derived from biochemical and biophysical studies with small molecule inhibitors.^17,20–22^ These exogenous ligands tap into a network of residues extending 20 Å away from the active site, which reorganize their noncovalent interactions in response to ligand binding to modulate catalysis. We hypothesize that this potential for allostery in PTP1B evolved to be engaged by endogenous ligands, such as proteins. Notably, some protein tyrosine phosphatases feature a catalytically dead pseudophosphatase domain that structurally interfaces with the catalytic domain at similar sites to those seen for allosteric small molecule ligands.^20,21,52^ In at least one such case, the pseudophosphatase domain enhances catalytic domain activity two-to three-fold, consistent with a mechanism comparable to allosteric activation by another protein.^53^ The potential for allostery is likely conserved across the protein tyrosine phosphatase family.^25^ Thus, the findings described in our study may be relevant to other members of the tyrosine phosphatase family, where regulation of function could also be accomplished through direct protein-protein interactions with the catalytic domain.

Many eukaryotic proteins have domains that can recognize proline-rich regions, raising the possibility that PTP1B may be allosterically regulated by a variety of binding partners. Indeed, our proteomics experiments suggest that many proteins other than Grb2 may also bind at or near the proline-rich region of PTP1B. We observed other SH3 domain-containing proteins in our proteomics dataset, but none that were enriched with PTP1B^WT^ over PTP1B^PRD^, suggesting they do not bind to the proline-rich region. A previous proteomics study in B cells identified several SH3 domain-containing proteins as interactors of PTP1B, but their binding sites were not determined.^14^ These interactors may be cell-type specific and therefore not observed in our study. Many of the hits in our proteomics analysis have not previously been connected to PTP1B signaling. These proteins highlight the potential of the proline-rich region as a regulatory handle for PTP1B function in new signaling contexts. One such example is the ER protein VAPA, discussed earlier. The signaling consequences of the PTP1B-VAPA interaction have not been defined, but our data strongly suggest a binding interface around the proline-rich region. The PRD mutation or other proximal mutations in PTP1B will likely be useful tools to untangle how PTP1B and VAPA cooperate to mediate focal adhesions. Whether VAPA or other interactors of the proline-rich region allosterically alter PTP1B activity remains to be seen.

The allosteric modulation of PTP1B by Grb2 may be impacted by other interactions and regulatory mechanisms controlling this phosphatase. This interaction is relatively weak (mid-micromolar), which means that other proteins could compete with Grb2 for occupancy of the proline-rich region. The relative local concentrations of these proteins will likely dictate signaling outcomes, and it may be the case that not every binder to the proline-rich region of PTP1B alters its catalytic activity. Thus, the proteins identified here as putative interactors of the proline-rich region, as well as SH3-containing proteins described by others,^13,14,24^ should be biochemically evaluated as putative regulators of PTP1B activity. Furthermore, PTP1B allosteric modulation by protein-protein interactions could be amplified or suppressed by other regulatory processes, such as enzymatic post-translational modifications, oxidation, or inhibitory protein-protein interactions.^37,48,54–56^ Thus, we anticipate that our findings will pave the way for new investigations into PTP1B regulation and signaling.

## Acknowledgements

We thank the members of the Shah and Keedy labs for their scientific insights and helpful discussions. We would also like to thank Lauren Tang for help with the proteomics experiments, as well as Blake Riley for his structural biology expertise and insights in the early stages of this work. We thank Rinat Abzalimov for assistance for HDX-MS experiments. We also thank Shibani Bhattacharya and Mike Goger of the New York Structural Biology Center (NYSBC) for their assistance with and resources for NMR experiments. This research was funded by NIH grant R35GM138014 to NHS, NIH grant R35GM133769 to DAK, and a Cottrell Scholar Award to DAK. CAC is supported by an NSF Graduate Research Fellowship (award #2036197). M.J. is funded by NIH grants R01AG071869 and R01HG012216, NSF award 2224211, and Columbia startup funding. The NMR experiments were supported by NIH NIGMS R01 GM088724 to AEM, and by The Center on Macromolecular Dynamics by NMR Spectroscopy (CoMD/NMR), a Biomedical Technology Development and Dissemination (BTDD) Center in turn supported by NIH RM1 GM145397.

## Supplementary Information

Materials and methods, along with all supplementary figures and references, are included in a single supplementary information file. Supplementary tables are provided as separate spreadsheets.

## Supplementary figures and tables

Figure S1. Grb2^WT^ binding to PTP1B^1-321^, PTP1B^PRD^, and PTP1B^1-401^.

Figure S2. Sortase labeling and fluorescence polarization measurements with Grb2 constructs.

Figure S3. Michaelis-Menten kinetics of PTP1B activity with phosphopeptides and *p*NPP.

Figure S4. Comparison of HDX-MS results and analysis of minor chemical shift changes observed by NMR.

Figure S5. Michaelis-Menten kinetics of PTP1B mutants with and without Grb2. Figure S6. Distribution of normalized intensities in MS proteomics replicates.

Table S1. NMR chemical shift assignments for apo PTP1B^1-321^ and PTP1B^1-321^ in complex with Grb2.

Table S2. MS proteomics data, including normalized intensities and MS2 counts for all protein groups, protein groups filtered by MS2 counts, fold-change values comparing PTP1B^WT^ to PTP1B^PRD^, and statistically significant hits.

