## Supplementary Information and Figures for "Allosteric regulation of the tyrosine phosphatase PTP1B by a protein-protein interaction"

**ORCiDs**

Cassandra A. Chartier: 0009-0004-2474-9937

Virgil A. Woods: 0000-0002-4796-8698

Yunyao Xu: 0000-0001-6587-9528

Anne E. van Vlimmeren: 0000-0003-0379-4945

Andrew C. Johns: 0000-0002-3197-1390

Marko Jovanovic: 0000-0001-9707-3377

Ann E. McDermott: 0000-0002-9249-1649

Daniel A. Keedy: 0000-0002-9184-7586

Neel H. Shah: 0000-0002-1186-0626

### This Supplementary Information file contains:

Materials and Methods pg 2

Figure S1. Grb2$^{WT}$ binding to PTP1B$^{1-321}$, PTP1B$^{PRD}$, and PTP1B$^{1-401}$. pg 10

Figure S2. Sortase labeling and fluorescence polarization measurements with Grb2 constructs. pg 11

Figure S3. Michaelis-Menten kinetics of PTP1B activity with phosphopeptides and pNPP. pg 12

Figure S4. Comparison of HDX-MS results and analysis of minor chemical shift changes observed by NMR. pg 13

Figure S5. Michaelis-Menten kinetics of PTP1B mutants with and without Grb2. pg 14

Figure S6. Distribution of normalized intensities in MS proteomics replicates. pg 15

Supplementary References pg 16

### Materials and Methods

#### Sequences of protein constructs used in this study

For each protein, the sequence of the full expression product is given, and the site for proteolytic removal of an affinity/expression tag is indicated with a vertical line. The part of the construct corresponding to native protein sequence is underlined. Mutations are shown in bold. All protein sequences used were derived from humans.

**PTP1B**

*Plasmid name:* pET-28b His$_6$-TEV-PTP1B cat
*Native residue range:* 1-321
*Protein sequence:*
MGSSHHHHHHDYDIPTTENLYFQ|GHMEMEKEFEQIDKSGSWAAIYQDIRHEASDFPCRVAKLPKNKN
RNRYRDVSPFDHSRIKLHQEDNDYINASLIKMEEAQRSYILTQGPLPNTCGHFWEMVWEQKSRGVVM
LNRVMEKGSLKCAQYWPQKEEKEMIFEDTNLKLTLISEDIKSYYTVRQLELENLTTQETREILHFHYTTW
PDFGVPESPASFLNFLFKVRESGSLSPEHGPVVVHCSAGIGRSGTFCLADTCLLLMDKRKDPSSVDIKK
VLLEMRKFRMGLIQTADQLRFSYLAVIEGAKFIMGDSSVQDQWKELSHEDLEPPPEHIPPPPRPPKRILE
PHN

*Plasmid name:* pET-28b His6-TEV-PTP1B cat PRD
*Native residue range:* 1-321
*Protein sequence:*
MGSSHHHHHHDYDIPTTENLYFQ|GHMEMEKEFEQIDKSGSWAAIYQDIRHEASDFPCRVAKLPKNKN
RNRYRDVSPFDHSRIKLHQEDNDYINASLIKMEEAQRSYILTQGPLPNTCGHFWEMVWEQKSRGVVM
LNRVMEKGSLKCAQYWPQKEEKEMIFEDTNLKLTLISEDIKSYYTVRQLELENLTTQETREILHFHYTTW
PDFGVPESPASFLNFLFKVRESGSLSPEHGPVVVHCSAGIGRSGTFCLADTCLLLMDKRKDPSSVDIKK
VLLEMRKFRMGLIQTADQLRFSYLAVIEGAKFIMGDSSVQDQWKELSHEDLE**AAA**EHI**AAAA**R**AA**KRIL
EPHN

*Note:* All biochemical experiments, HDX-MS, and NMR were performed using the 1-321 native residue range.

*Plasmid name:* pEF Myc-PTP1B FL WT
*Native residue range:* 1-435
*Protein sequence:*
MEQKLISEEDLGGQSGQMEMEKEFEQIDKSGSWAAIYQDIRHEASDFPCRVAKLPKNKNRNRYRDVS
PFDHSRIKLHQEDNDYINASLIKMEEAQRSYILTQGPLPNTCGHFWEMVWEQKSRGVVMLNRVMEKG
SLKCAQYWPQKEEKEMIFEDTNLKLTLISEDIKSYYTVRQLELENLTTQETREILHFHYTTWPDFGVPES
PASFLNFLFKVRESGSLSPEHGPVVVHCSAGIGRSGTFCLADTCLLLMDKRKDPSSVDIKKVLLEMRKF
RMGLIQTADQLRFSYLAVIEGAKFIMGDSSVQDQWKELSHEDLEPPPEHIPPPPRPPKRILEPHNGKCR
EFFPNHQWVKEETQEDKDCPIKEEKGSPLNAAPYGIESMSQDTEVRSRVVGGSLRGAQAASPAKGEP
SLPEKDEDHALSYWKPFLVNMCVATVLTAGAYLCYRFLFNSNT

*Plasmid name:* pEF Myc-PTP1B FL PRD
*Native residue range:* 1-435
*Protein sequence:*
MEQKLISEEDLGGQSGQMEMEKEFEQIDKSGSWAAIYQDIRHEASDFPCRVAKLPKNKNRNRYRDVS
PFDHSRIKLHQEDNDYINASLIKMEEAQRSYILTQGPLPNTCGHFWEMVWEQKSRGVVMLNRVMEKG
SLKCAQYWPQKEEKEMIFEDTNLKLTLISEDIKSYYTVRQLELENLTTQETREILHFHYTTWPDFGVPES
PASFLNFLFKVRESGSLSPEHGPVVVHCSAGIGRSGTFCLADTCLLLMDKRKDPSSVDIKKVLLEMRKF
RMGLIQTADQLRFSYLAVIEGAKFIMGDSSVQDQWKELSHEDLE**AAA**EHI**AAAA**R**AA**KRILEPHNGKC
REFFPNHQWVKEETQEDKDCPIKEEKGSPLNAAPYGIESMSQDTEVRSRVVGGSLRGAQAASPAKGE
PSLPEKDEDHALSYWKPFLVNMCVATVLTAGAYLCYRFLFNSNT

*Note:* Cell biology experiments were performed using the full-length 1-435 native residue range.

**Grb2**

*Plasmid name:* pET-28b His6-TEV-Grb2 WT
*Native residue range:* 1-217
*Protein sequence:*
MGSSHHHHHHDYDIPTTENLYFQ|GHMASMTGGQQMGRGSMEAIAKYDFKATADDELSFKRGDILKVL
NEECDQNWYKAELNGKDGFIPKNYIEMKPHPWFFGKIPRAKAEEMLSKQRHDGAFLIRESESAPGDFS
LSVKFGNDVQHFKVLRDGAGKYFLWVVKFNSLNELVDYHRSTSVSRNQQIFLRDIEQVPQQPTYVQAL
FDFDPQEDGELGFRRGDFIHVMDNSDPNWWKGACHGQTGMFPRNYVTPVNRNV

*Plasmid name:* pET-28b His6-TEV-Grb2 Y160E
*Native residue range:* 1-217
*Protein sequence:*
MGSSHHHHHHDYDIPTTENLYFQ|GHMASMTGGQQMGRGSMEAIAKYDFKATADDELSFKRGDILKVL
NEECDQNWYKAELNGKDGFIPKNYIEMKPHPWFFGKIPRAKAEEMLSKQRHDGAFLIRESESAPGDFS
LSVKFGNDVQHFKVLRDGAGKYFLWVVKFNSLNELVDYHRSTSVSRNQQIFLRDIEQVPQQPT**E**VQAL
FDFDPQEDGELGFRRGDFIHVMDNSDPNWWKGACHGQTGMFPRNYVTPVNRNV

*Plasmid name:* pET His6-SUMO-Grb2 N-SH3
*Native residue range:* 1-58
*Protein sequence:*
MGSSHHHHHHGSGLVPRGSASMSDSEVNQEAKPEVKPEVKPETHINLKVSDGSSEIFFKIKKTTPLRR
LMEAFAKRQGKEMDSLRFLYDGIRIQADQTPEDLDMEDNDIIEAHREQIGG|MEAIAKYDFKATADDELS
FKRGDILKVLNEECDQNWYKAELNGKDGFIPKNYIEMKPH

*Plasmid name:* pET His6-SUMO-Grb2 SH2
*Native residue range:* 60-152
*Protein sequence:*
MGSSHHHHHHGSGLVPRGSASMSDSEVNQEAKPEVKPEVKPETHINLKVSDGSSEIFFKIKKTTPLRR
LMEAFAKRQGKEMDSLRFLYDGIRIQADQTPEDLDMEDNDIIEAHREQIGG|WFFGKIPRAKAEEMLSK
QRHDGAFLIRESESAPGDFSLSVKFGNDVQHFKVLRDGAGKYFLWVVKFNSLNELVDYHRSTSVSRN
QQIFLRDIE

*Plasmid name:* pET His6-SUMO-Grb2 C-SH3
*Native residue range:* 156-215
*Protein sequence:*
MGSSHHHHHHGSGLVPRGSASMSDSEVNQEAKPEVKPEVKPETHINLKVSDGSSEIFFKIKKTTPLRR
LMEAFAKRQGKEMDSLRFLYDGIRIQADQTPEDLDMEDNDIIEAHREQIGG|QQPTYVQALFDFDPQED
GELGFRRGDFIHVMDNSDPNWWKGACHGQTGMFPRNYVTPVNR

*Note:* All individual domain constructs used for sortase-mediated labeling were cloned with an additional two glycine residues before the beginning of the native protein sequence.

### Expression and Purification of Proteins

For over-expression, BL21(DE3) cells were transformed with plasmids encoding each protein. Cells were grown in terrific broth (TB) supplemented with 50 µg/mL kanamycin at 37 °C until cells reached an optical density at 600 nm ($OD_{600}$) of 0.5. IPTG (0.5 mM) was added to induce protein expression. After induction, protein expression was carried out at 18 °C overnight. Cells were pelleted by centrifugation at 4000 RCF and subsequently resuspended in lysis buffer (50 mM Tris pH 8.0, 300 mM NaCl, 10 mM imidazole, 10% glycerol, and freshly added 2 mM β-mercaptoethanol). The cells were lysed using sonication (Fisherbrand Sonic Dismembrator), and spun down at 42,587 RCF for 45 minutes. The supernatant was applied to a 5 mL HisTrap HP column (Cytiva). The resin was washed with 10 column volumes of lysis buffer and 10 column volumes of wash buffer (50 mM Tris pH 8.5, 50 mM NaCl, 10 mM imidazole, 10% glycerol, and freshly added 2 mM β-mercaptoethanol). The protein was eluted off the HisTrap HP column in using 250 mM imidazole and brought onto a 5mL HiTrap Q anion exchange column (Cytiva). The column was washed using Anion A buffer, where the pH depended on the isoelectric point of each protein (50 mM Tris, 50 mM NaCl, 10% glycerol, and 1 mM TCEP). Protein elution off the column was induced through a salt gradient between Anion A buffer and Anion B buffer (50 mM Tris pH, 1 M NaCl, 1 mM TCEP, and 10% glycerol). The eluted protein was cleaved at the respective proteolytic cleavage site by addition of 0.05 mg/mL $His_6$-tagged protease at 4 °C overnight. This cleavage cocktail was flowed through a 5 mL HisTrap HP column (Cytiva) to isolate the cleaved protein away from uncleaved protein and the respective protease. Finally, the cleaved protein was purified by size-exclusion chromatography on a gel filtration column (Cytiva) equilibrated with SEC buffer (10 mM HEPES pH 7.5, 150 mM NaCl, 1mM TCEP, and 10% glycerol). Pure fractions were pooled and concentrated, and flash frozen in liquid $N_2$ for long-term storage at −80 °C.

### Purification of PTP1B

All PTP1B constructs for bacterial expression were cloned into a kanamycin-resistant pET plasmid encoding an N-terminal $His_6$-tag and TEV protease cleavage site. Anion exchange chromatography was performed using Anion A/B buffers at pH 8.5. The $His_6$-TEV tag was cleaved using TEV protease. The cleaved protein was purified using a Superdex 75 10/300 column.

### Purification of Grb2

All full-length Grb2 variants for bacterial expression were cloned into a kanamycin-resistant pET plasmid encoding an N-terminal $His_6$-tag and TEV protease cleavage site. Anion exchange chromatography was performed using Anion A/B buffers at pH 7.5. The $His_6$-TEV tag was cleaved using TEV protease. Cleaved $Grb2^{WT}$ was purified using a Superdex 200 16/600 column, while $Grb2^{Y160E}$ was purified using a Superdex 200 10/300 column.

### Purification of Grb2 SH3 domains

Grb2 N-SH3 and C-SH3 were cloned into $His_6$-SUMO-N-SH3 and $His_6$-SUMO-C-SH3 constructs, respectively. Anion exchange chromatography was performed using Anion A/B buffers at pH 6.5. The $His_6$-SUMO-TEV tag was cleaved using Ulp1 protease. The cleaved protein was purified using a Superdex 75 10/300 column.

### Purification of Grb2 SH2 domain

Grb2 SH2 was cloned into a $His_6$-SUMO-SH2 construct. Anion exchange chromatography was performed using Anion A/B buffers at pH 10.5. The $His_6$-TEV tag was cleaved using Ulp1 protease. The cleaved protein was purified using a Superdex 75 10/300 column.

### Size exclusion chromatography binding assays

PTP1B and Grb2 variants were thawed at room temperature. 20 µM samples of PTP1B and Grb2 variants alone as well as an equimolar (20 µM each) mixture of the two proteins were prepared in SEC buffer (10 mM HEPES pH 7.5, 150 mM NaCl, 1mM TCEP, and 10% glycerol). The samples were then injected onto a Superdex 200 10/300 gel filtration column (Cytiva) equilibrated with SEC buffer and absorbance was monitored at 280 nm. Fractions were collected for SDS-PAGE analysis.

### Synthesis and purification of fluorescent peptide for sortase-mediated labeling

The fluorescent peptide (FITC-Ahx-LPETGG-$NH_2$) was synthesized using 9-fluorenylmethoxycarbonyl (Fmoc) solid-phase peptide chemistry. Synthesis was carried out using the Liberty Blue automated microwave-assisted peptide synthesizer from CEM under nitrogen atmosphere, with standard manufacturer-recommended protocols. The peptide was synthesized on MBHA Rink amide resin solid support (0.1 mmol scale). Each Nα-Fmoc amino acid (6 eq, 0.2 M) was activated with diisopropylcarbodiimide (DIC, 1.0 M) and ethyl cyano(hydroxyamino)acetate (Oxyma Pure, 1.0 M) in dimethylformamide (DMF) prior to coupling. All Nα-Fmoc amino acid coupling cycles were done at 75 °C for 15 s, then 90 °C for 110 s. Deprotection of the Fmoc group was performed in 20% (v/v) piperidine in DMF (75 °C for 15 s then 90 °C for 50 s). The resin was washed (4x) with DMF following Fmoc deprotection and after Nα-Fmoc amino acid coupling. Then, Fmoc-Ahx (6 eq, 0.2 M) was activated with diisopropylcarbodiimide (DIC, 1.0 M) and ethyl cyano(hydroxyamino)acetate (Oxyma Pure, 1.0 M) in dimethylformamide (DMF) prior to coupling. The coupling cycle was done at 75 °C for 35 s then 90 °C for 575 s. Removal of the terminal Fmoc group was performed as described above.

After peptide synthesis was completed, including N-terminal Ahx attachment and deprotection, the resin was washed (3x each) with dichloromethane (DCM) and methanol (MeOH) and dried under reduced pressure overnight. Then, a portion of the resin (0.025 mmol) was prepared for FITC labeling by swelling in DMF with agitation for 30 min. Excess DMF was removed and to the resin was added fluorescein isothiocyanate (FITC) (0.075 mmol, 3 eq.) and DIPEA (0.15mmol, 6 eq.) in DMF. This reaction was incubated at room temperature with agitation for 2 hours. After FITC labeling, the resin was washed (3x each) with dichloromethane (DCM) and methanol (MeOH) and dried under reduced pressure overnight.

The crude fluorescent peptide mixture was cleaved and purified using reverse-phase high performance liquid chromatography (RP-HPLC) on a semi-preparatory C18 column (Agilent, ZORBAX 300SB-C18, 9.4 x 250 mm, 5 µm) with an Agilent HPLC system (1260 Infinity II). Flow rate for purification was kept at 4 mL/min with solvents A (water, 0.1% (v/v) TFA) and B (acetonitrile, 0.1% (v/v) TFA). The peptide was purified over a 20-30% B gradient in 40 minutes. Peptide purity was assessed with an analytical column (Agilent, ZORBAX 300SB-C18, 4.6 x 150 mm, 5 µm) at a flow rate of 1 mL/min over a 0-70% B gradient in 30 minutes. The peptide was determined to be ≥95% pure by peak integration and its identity was confirmed by mass spectroscopy (Waters Xevo G2-XS QTOF). The pure peptide was lyophilized and redissolved in PBS pH 8.0 as needed for experiments.

### Sortase-mediated labeling of isolated Grb2 domains

Each isolated Grb2 domain with two N-terminal glycines was diluted to a concentration of 76.2 µM in reaction buffer (50mM Tris pH 7.5, 150mM NaCl) with 2.6 µM sortase 7M[1] (prepared from Addgene plasmid #51141) and 400 µM fluorescently-labeled peptide. The mixture was incubated at 37 °C for 3 hours. The reaction mixture was diluted with lysis buffer (50 mM Tris pH 8.0, 300 mM NaCl, 10 mM imidazole, 10% glycerol, and freshly added 2 mM β-mercaptoethanol) and applied to a 1 mL Ni-NTA column (Cytiva) to to isolate the fluorescently-labeled protein away from the sortase. Finally, the fluorescently-labeled protein was further purified by size-exclusion chromatography on a Superdex 75 10/300 gel filtration column (Cytiva) equilibrated with SEC buffer (10 mM HEPES pH 7.5, 150 mM NaCl, 1mM TCEP, and 10% glycerol). Pure fractions were pooled and concentrated, and flash frozen in liquid $N_2$ for long-term storage at −80 °C.

### Fluorescence anisotropy binding assays

Each fluorescently-labeled Grb2 isolated domain and PTP1B constructs were thawed in room temperature water. PTP1B constructs were first concentrated to ~1.1 mM using a centrifugal concentrator and then buffer exchanged with assay buffer using a centrifugal concentrator (60mM HEPES pH 7.2, 75 mM KCl, 75 mM NaCl,1 mM EDTA, 0.05% Tween-20). They were then serially diluted 13 times in assay buffer, with a 1.1x starting concentration. Each fluorescently-labeled Grb2 domain was diluted to 10X the desired concentration and transferred to a black 96-well plate. Each PTP1B construct titration was then transferred to the well plate and mixed in a 10:1 ratio with the fluorescently-labeled Grb2 domain. The mixture was incubated for 15 minutes at room temperature. Parallel and perpendicular measurements were taken using the 485/30 polarization cube on the BioTek Neo2 Plate Reader. Data was analyzed and fitted to a quadratic binding equation to determine the $K_D$ for each fluorescently-labeled Grb2 domain, according to previously established methods.[2,3]

### Synthesis and purification of phosphopeptides for activity measurements

Phosphopeptides were also synthesized using 9-fluorenylmethoxycarbonyl (Fmoc) solid-phase peptide chemistry as described above for the fluorescent peptide. The coupling cycles for phosphotyrosine and the amino acid directly after it were done at 75 °C for 15 s, then 90 °C for 230 s. All other coupling cycles were done at 75 °C for 15 s, then 90 °C for 110 s. Deprotection of the Fmoc group was performed in 20% (v/v) piperidine in DMF (75 °C for 15 s then 90 °C for 50 s), except for the amino acid directly after the phosphotyrosine which had an additional initial deprotection (25 °C for 300 s). The resin was washed (4x) with DMF following Fmoc deprotection and after Nα-Fmoc amino acid coupling. All peptides were acetylated at their N-terminus with 10% (v/v) acetic anhydride in DMF and washed (4x) with DMF.

After peptide synthesis was completed, including N-terminal acetylation, the resin was washed (3x each) with dichloromethane (DCM) and methanol (MeOH), and dried under reduced pressure overnight. The peptides were cleaved and the side chain protecting groups were simultaneously deprotected in 95% (v/v) trifluoroacetic acid (TFA), 2.5% (v/v) triisopropylsilane (TIPS), and 2.5% water, in a ratio of 10 µL cleavage cocktail per mg of resin. The cleavage-resin mixture was incubated at room temperature for 90 minutes, with agitation. The cleaved peptides were precipitated in cold diethyl ether, washed in ether, pelleted, and dried under air. The peptides were redissolved in a 50% (v/v) water/acetonitrile solution and filtered from the resin.

Each crude peptide mixture was purified using reverse-phase high performance liquid chromatography (RP-HPLC) on either a semi-preparatory C18 column (Agilent, ZORBAX 300SB-C18, 9.4 x 250 mm, 5 µm) with an Agilent HPLC system (1260 Infinity II), or a preparatory C18 column (XBridge Peptide BEH C18 Prep Column, 19 x 150 mm, 5 µm) with a Waters prep-HPLC system (Prep 150 LC System). Flow rate for purification was kept at 4 mL/min (semi-preparative) or 17mL/min (preparative) with solvents A (water, 0.1% (v/v) TFA) and B (acetonitrile, 0.1% (v/v) TFA). Peptides were generally purified over a 40 minute (semi-preparative) or 13 minute (preparative) linear gradient from solvent A to solvent B, with the specific gradient depending on the peptide sample. Peptide purity and identity were assessed as described above for the fluorescent peptide. Pure peptides were lyophilized and redissolved in 100 mM Tris, pH 8.0, as needed for experiments.

### PTP1B activity measurements

Initial rate measurements for the PTP1B-catalyzed dephosphorylation of *p*-nitrophenyl phosphate (pNPP) and various phosphopeptide substrates were conducted at room temperature in buffer (10 mM HEPES pH 7.5, 150 mM NaCl, 1mM TCEP, and 10% glycerol). For pNPP, reactions of 200 µL were set up in a clear polystyrene flat bottom 96-well plate. A substrate concentration series of 0-6 mM was used to generate Michaelis-Menten curves. Reactions were started by the addition of 100 nM PTP1B construct and 50 µM $Grb2^{WT}$. Activity measurements using phosphopeptides were done using the EnzCheck Phosphate Assay Kit (ThermoFisher) according to the manufacturer's instructions. Reactions of 100 µL were set up in a clear polystyrene flat bottom half area 96-well plate. A substrate concentration series of 0-1000 µM or 0-500 µM was used to determine $k_{cat}$ and $K_M$. Reactions were started by addition of 20 nM PTP1B construct and 50 µM $Grb2^{Y160E}$. For the $Grb^{Y160E}$ and isolated C-SH3 domain titration experiments, centrifugal concentrators were used to achieve a concentration series of 0-900 µM and 0-1000 µM, respectively. Absorbance at 360 nm was recorded every 3-4 seconds in a span of 2 minutes using a BioTek Synergy Neo2 multi-mode reader.

In all cases, the linear region of the reaction progress curve was determined by visual inspection and fit to a line. These slopes were converted from absorbance units as a function of time to product formation as a function of time using standard curves measured with inorganic phosphate. Finally, these rates were corrected for enzyme concentration used in the experiment to yield $V_0$ / [enzyme] in units of ($s^{-1}$). These corrected rates were plotted as a function of substrate concentration and fit to the Michaelis-Menten equation using non-linear regression to determine catalytic parameters. Experiments were generally repeated at least three times, and the average and standard deviation of all individual replicates are reported.

### NMR spectroscopy

NMR data were collected on 800 MHz spectrometers equipped with Bruker Advance II console and 5 mm CTI cryoprobe. NMR measurements of $PTP1B^{1-321}$ with and without $Grb2^{WT}$ were recorded using $^{15}N$-labeled $PTP1B^{1-321}$ at a final concentration of 0.15 mM in NMR buffer (50mM HEPES, pH 6.8, 150mM NaCl, 1mM TCEP in 5% D2O). The sequence-specific backbone assignments of PTP1B were assigned by comparison to previously published spectra (BMRB ID: 19224).[4] All NMR data were analyzed using CcpNMR.[5]

### Hydrogen-deuterium exchange mass spectrometry

*Sample handling*

For intact mass HDX experiments, all sample handling was performed using the LEAP HDX platform. This automated system initiates and precisely times the labeling reactions (reaction volume of 50 µL), followed by rapid mixing with a quench solution to reduce the temperature to 0-4°C. The quenched samples were then immediately flowed into the mass spectrometer for analysis.

*HDX labeling*

Purified apo protein constructs ($PTP1B^{WT}$ and $PTP1B^{PRD}$) were prepared in a buffer containing 10 mM HEPES pH 7.5, 150 mM NaCl, and 1 mM TCEP. For experiments involving the $Grb2^{Y160E}$ and the isolated N- and C-SH3 domains, the protein samples were incubated with varying concentrations of the $Grb2^{Y160E}$ complex or isolated N- and C-SH3 domains. All HDX reactions were carried out at 15 °C.

The $PTP1B^{WT}$ and $PTP1B^{PRD}$ protein samples were mixed with one of the three Grb2 constructs before labeling at concentrations 5x the nominal concentrations shown in **Figure 4B**. For labeling, the protein was mixed with a $D_2O$ buffer identical in composition to the $H_2O$ buffer but prepared from scratch using $D_2O$, achieving a final $D_2O/H_2O$ purity of >99.5%. The mixed protein preparations were diluted (1:5) in the $D_2O$ buffer for a final $D_2O$ content of 80% in the reaction mixture. The labeling reaction was conducted for different time points in replicate experiments: 30 s (n=3), 300 s (n=2), and 3000 s (n=2). HDX quenching was performed by mixing the labeling reaction with a cold quenching solution (1.5% formic acid) in a 1:1 ratio, halting the HDX reaction.

*Mass spectrometry analysis*

The quenched samples were injected into a Bruker maXis-II ESI-QqTOF high-resolution mass spectrometer for intact mass analysis. This setup allowed for direct measurement of the intact protein mass, providing a global overview of the deuterium incorporation across all protein molecules in the sample.

*Preprocessing and data alignment*

Raw mass spectrometry data files were processed using Compass Data Analysis 5.3. The centroid m/z value of intact $PTP1B^{WT}$ or $PTP1B^{PRD}$ were calculated by weighted average of the peaks identified. Data alignment was performed based on protein mass, retention time, and m/z values to ensure accurate comparison across different time points.

*Deuteration level calculation*

The deuteration level of the intact protein was determined by comparing the centroid mass of the deuterated protein to the non-deuterated protein at each time point. This comparison provided the deuteration percentage over time, reflecting the overall dynamics and structural changes of the protein.

### Cell culture and cell biological experiments

*General cell culture information*

Human embryonic kidney (HEK) 293 cells were cultured in Dulbecco's Modified Eagle Medium (DMEM) supplemented with 10% Fetal Bovine Serum (FBS) and 1% penicillin/streptomycin at 37 °C with 5% $CO_2$.

*DNA constructs*

The full-length PTP1B gene was cloned from the pPTP1BD181A-mCherry plasmid (Addgene plasmid #40270). The proline residues in the proline-rich region were mutated to alanine to generate a proline-rich dead (PRD) PTP1B mutant. These genes, full-length $PTP1B^{WT}$ and $PTP1B^{PRD}$, were cloned into a pEF vector (a gift from the Arthur Weiss lab) for transient transfection.

*Co-immunoprecipitation experiments*

~7 x $10^6$ HEK 293 cells were seeded in a 15 cm plate. The next day, the cells were transfected using 17 µg and 25 µg of $PTP1B^{WT}$ and $PTP1B^{PRD}$ plasmids, respectively, to ensure equal expression. The plasmids were added to 2.5 mL DMEM, followed by the addition of 51 µg and 75 µg PEI to the $PTP1B^{WT}$ and $PTP1B^{PRD}$ plasmid mixtures, respectively. The transfection medium was refreshed after 16 hours and replaced with complete medium. The cells were harvested 48 hours after transfection by scraping in PBS. The cells were washed once in PBS and lysed in 400 µL lysis buffer (20mM Tris, pH 8.0, 137mM NaCl, 10% glycerol, 1% Triton X-100, 2mM EDTA + protease inhibitors + phosphatase inhibitors) for 25 minutes while rotating at 4 °C. The insoluble material was pelleted using centrifugation at 54,036 RCF for 15 minutes at 4 °C. The supernatant was transferred to a clean microcentrifuge tube and stored at -20 °C.

A bicinchoninic acid (BCA) assay was used to determine protein concentration by measuring absorbance at 562 nm using a BioTek Synergy Neo2 multi-mode reader. 1000 µg of protein in a total volume of 400 µL was incubated overnight at 4 °C with 25 µg of magnetic Myc-beads while rotating. The next day, the beads were washed 3X with 1 mL lysis buffer using a magnetic rack. Then, 97.5 µL 1X Laemmli buffer was added to the beads and they were boiled at 100 °C for 5 minutes. For whole cell lysates, 15 µg protein was loaded onto a gel. For IP samples, 15 µL of boiled supernatant was used. The gel was transferred onto a nitrocellulose membrane using TurboBlot (BioRad) and the membrane was blocked using 5% bovine serum albumin (BSA) in Tris-buffered saline (TBS) for 1 hour at room temperature. Membranes were rinsed with TBS with 0.1% Tween-20 (TBS-T) and incubated with primary antibodies in TBST + 5% BSA overnight at 4ºC (Myc 1:1000, GAPDH 1:1000, Grb2 1:500, ARHGAP12 1:500). Membranes were washed and incubated with secondary antibodies (IRDye 680 and 800). Blots were imaged on an Amersham Typhoon 5 Biomolecular Imager.

### Mass spectrometry proteomics

*Sample preparation*

The co-immunoprecipitation was done as described above, except with 2000 µg of protein. After overnight incubation at 4 °C, the beads were washed twice with 1 mL 50 mM Tris pH 8.0 and 200 µL 2 M urea in 50 mM Tris pH 8.0. After washing, the beads were resuspended in 80 µL of 2M urea in 50 mM

Tris pH 8.0 with 1 mM DTT and 0.4 µg trypsin for 1 hour at room temperature with agitation. The supernatants were then transferred to a clean microcentrifuge tubes. The beads were washed twice with 60 µL of 2M urea in 50 mM Tris pH 8.0, and the washes were combined with the supernatants. 4 mM DTT was added to each sample and incubated for 30 minutes at room temperature with agitation. Then, 10 mM iodoacetamide was added to each sample and incubated in the dark for 45 minutes at room temperature with agitation. An additional 0.5 µg trypsin was then added to each sample and incubated in the dark overnight at room temperature with agitation. The samples were then acidified using ~1% (v/v) formic acid (FA) and desalted using C18 stage tips. Before desalting, tips were conditioned with 100 µL MeOH, 100 µL 0.2% FA/60% ACN, and twice with 100 µL 0.2% FA. The samples were then loaded onto the stage tips and washed twice with 100 µL 0.2% FA. Finally, the peptides from each sample were eluted from the stage tips using 50 µL 0.2% FA/60% ACN and dried using a vacuum centrifuge. The peptides were stored in the -80 °C until resuspension and injection.

*LC-MS/MS analysis on a Q-Exactive HF*

Samples were resuspended in 13 µL 3%Acn/0.2%FA. 6 µL was injected and analyzed on a Waters M-Class UPLC using a 25cm Ionopticks Aurora column coupled to a benchtop Thermo Fisher Scientific Orbitrap Q Exactive HF mass spectrometer. Peptides were separated at a flow rate of 400 nL/min with a 100 min gradient, including sample loading and column equilibration times. Data was acquired in data dependent mode using Xcalibur 4.1 software. MS1 Spectra were measured with a resolution of 120,000, an AGC target of 3e6 and a mass range from 300 to 1800 m/z. Up to 12 MS2 spectra per duty cycle were triggered at a resolution of 60,000, an AGC target of 1e5, an isolation window of 0.8 m/z, a normalized collision energy of 28, a scan range of 200 to 2000 m/z, and a fixed first mass of 110 m/z.

*Quantification and statistical analyses*

All raw data were analyzed with SpectroMine software version 4.2.230428 using a UniProt database (Homo sapiens, UP000005640). Carbamidomethylation on cysteines was set as a fixed modification. Oxidation of methionine and protein N-terminal acetylation were set as variable modifications, with a maximum of 5 variable modifications. Trypsin/P was set as the digestion enzyme, and up to two missed cleavages were permitted. For identification, we applied a maximum false discovery rate of 1% on protein and peptide level. We required 1 or more unique or razor peptides for protein identification.

Common contaminants (such as keratins) were removed from further analysis. Then, noise was added from the randomly sampled lower range of the limit of detection to all raw intensities. Protein groups with <4.99 average MS/MS counts were then removed from further analysis. Protein group intensities were normalized for the total intensity of all observable protein groups in that sample. Normalized protein group intensities were log2-transformed and averaged, from which fold-changes were calculated. P-values were calculated using a two-tailed, heteroscedastic t-test.

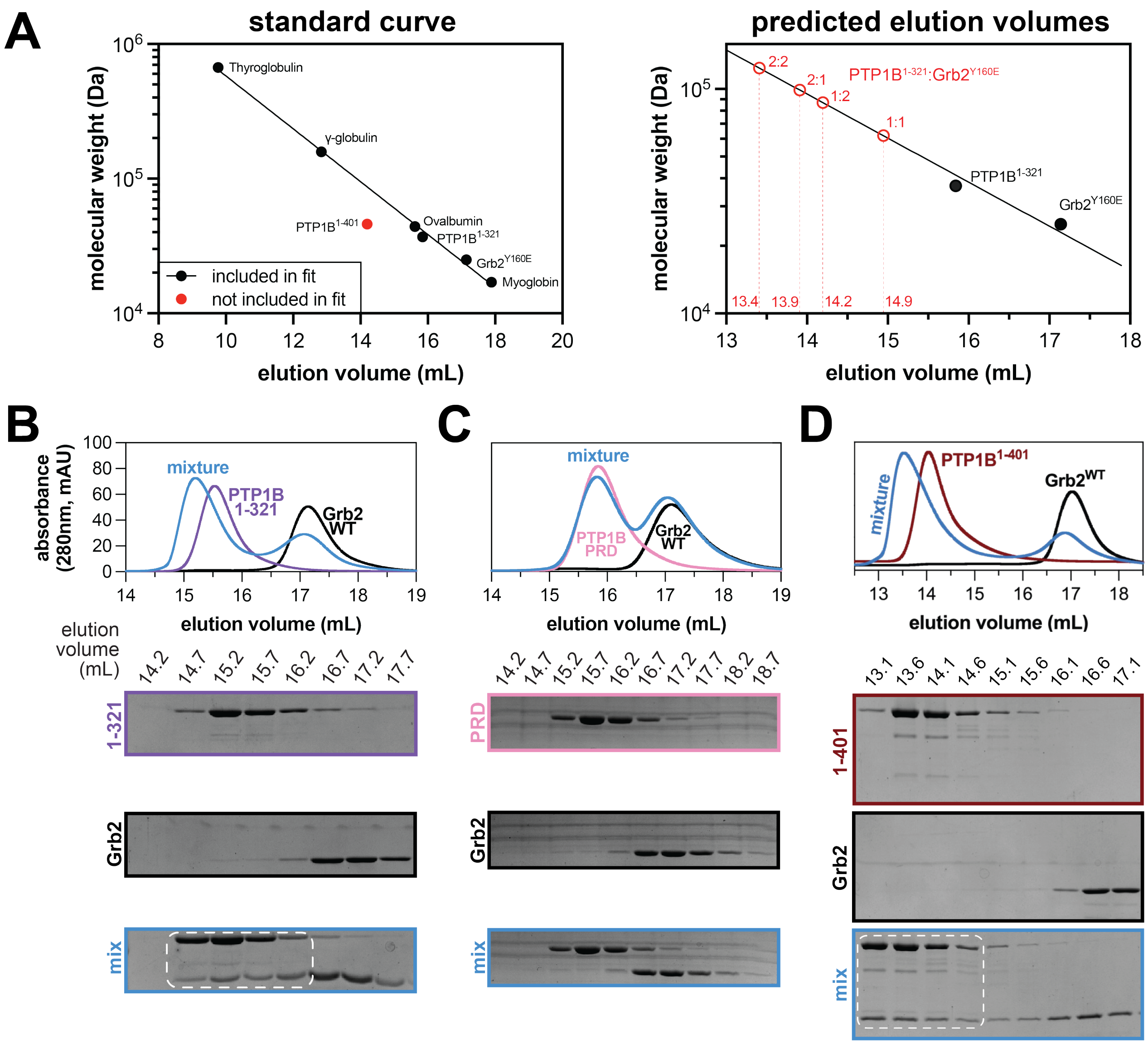


**Figure S1. Grb2$^{WT}$ binding to PTP1B$^{1\text{-}321}$, PTP1B$^{PRD}$, and PTP1B$^{1\text{-}401}$.** (**A**) Left panel: Size exclusion chromatography standard curve line relating $\log_{10}$ of protein molecular weight to elution volume, using PTP1B$^{1\text{-}321}$, Grb2$^{Y160E}$, and four unrelated globular proteins. Note that PTP1B$^{1\text{-}401}$ falls off of the standard curve because it has a large disordered region, thus it was not included in the fit. Right panel: predicted molecular weights for PTP1B$^{1\text{-}321}$:Grb2$^{Y160E}$ complexes of different stoichiometry, based on the standard curve line shown in the left panel. The 1:1 complex has an elution volume consistent with elution of the complex, as seen in **Figure 2A**. (**B**) Size exclusion chromatograms (top) depicting separate injections of equimolar PTP1B$^{1\text{-}321}$, Grb2$^{WT}$, and a mixture of the two proteins. SDS-PAGE analysis (bottom) shows co-elution of PTP1B$^{1\text{-}321}$ and Grb2$^{WT}$ at earlier elution volumes in the mixture. (**C**) Size exclusion chromatograms (top) depicting separate injections of equimolar PTP1B$^{PRD}$, Grb2$^{WT}$, and a mixture of the two proteins. SDS-PAGE analysis (bottom) shows the proteins in the mixture elute at the same volumes as when not in a mixture. (**D**) Size exclusion chromatograms (top) depicting separate injections of equimolar PTP1B$^{1\text{-}401}$, Grb2$^{WT}$, and a mixture of the two proteins. SDS-PAGE analysis (bottom) shows co-elution of PTP1B$^{1\text{-}401}$ and Grb2$^{WT}$ at earlier elution volumes in the mixture.

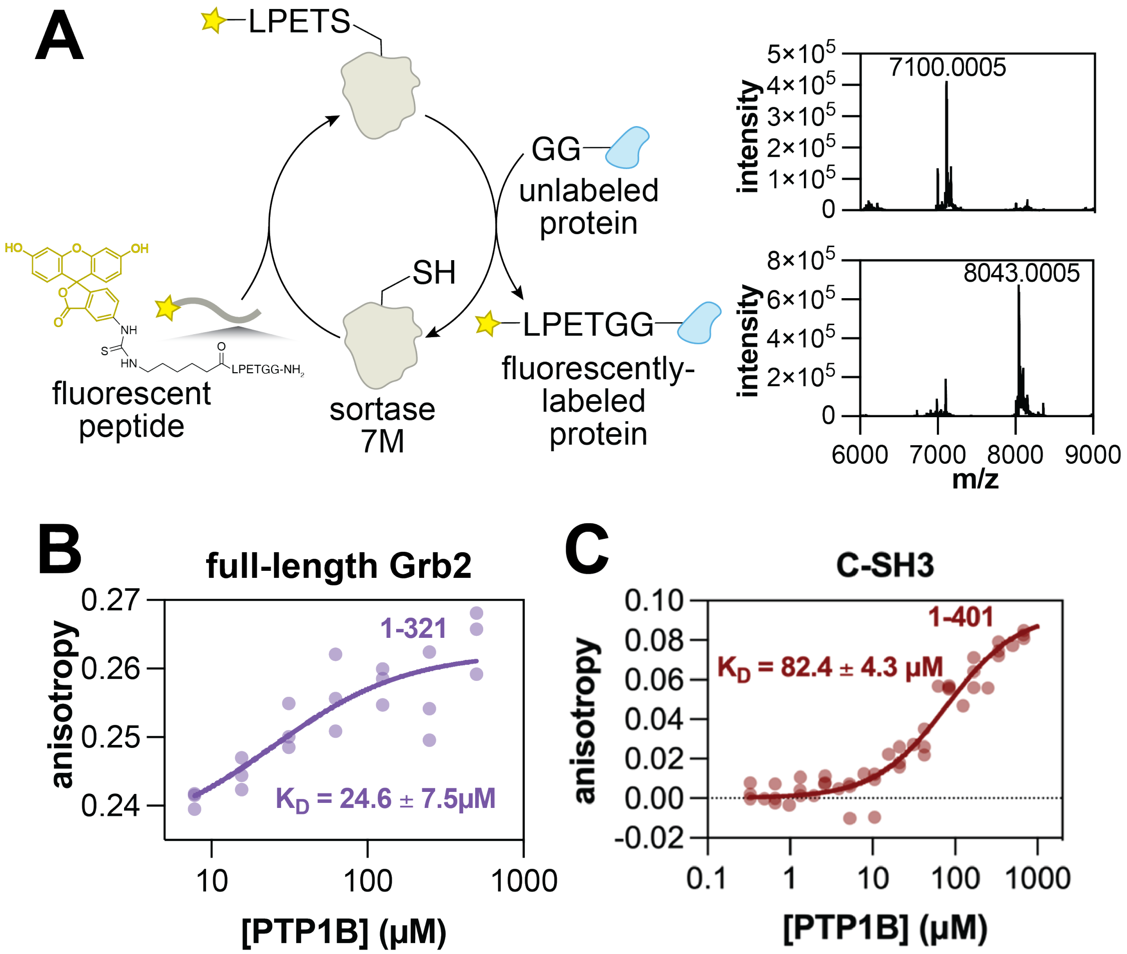


**Figure S2. Sortase labeling and fluorescence polarization measurements with Grb2 constructs.** (**A**) Fluorescent labeling of Grb2 constructs is achieved by sortase-mediated transpeptidation of a fluorescently-labeled peptide to a Grb2 construct with two N-terminal glycine residues. Expected masses for unlabeled protein and fluorescently labeled protein were 7101.8 Da and 8044.8 Da, respectively. Deconvoluted mass spectra for the fluorescently-labeled Grb2 C-SH3 domain is shown as an example. (**B**) Fluorescence anisotropy analysis of PTP1B$^{1\text{-}321}$ binding to fluorescently-labeled full-length Grb2$^{WT}$. (**C**) Fluorescence anisotropy analysis of PTP1B$^{1\text{-}401}$ binding to fluorescently-labeled Grb2 C-SH3 domain.

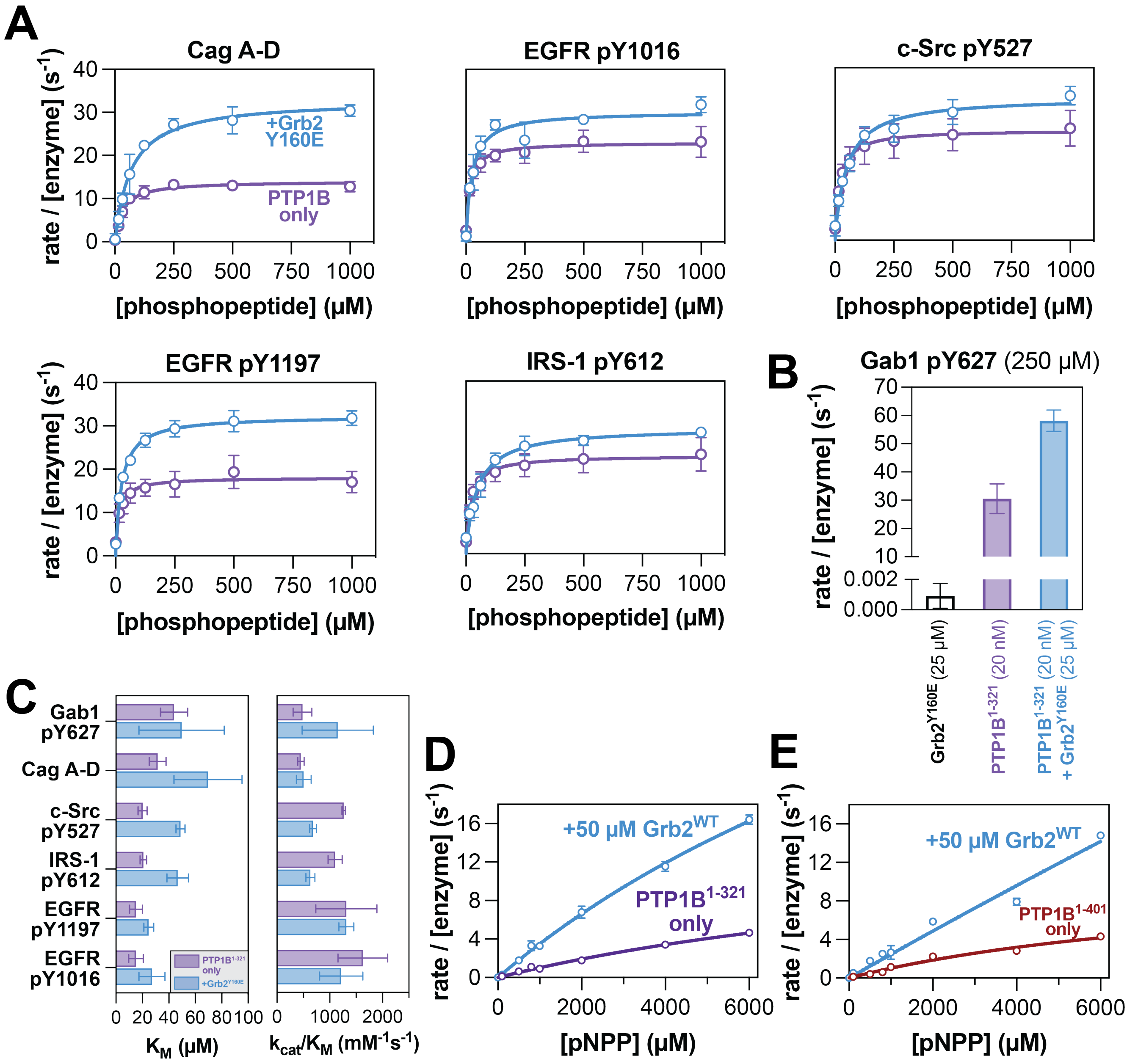


**Figure S3. Michaelis-Menten kinetics of PTP1B activity with phosphopeptides and *p*NPP.** (**A**) Michaelis-Menten curves showing PTP1B$^{1\text{-}321}$ activity alone (purple) and in response to 50 µM Grb2 Y160E (blue) against the rest of the phosphopeptide substrates. (**B**) Control experiment showing a lack of Gab1 pY627 dephosphorylation by 25 µM Grb2$^{Y160E}$ alone, compared with robust dephosphorylation by 20 nM PTP1B$^{1\text{-}321}$ alone and further rate enhancement with 20 nM PTP1B$^{1\text{-}321}$ + 25 µM Grb2$^{Y160E}$. Note that all of the observed rates in units of M/s were divided by the PTP1B concentration used in the latter two measurements (20 nM) for direct comparison. Thus, the absolute signal for 25 µM Grb2$^{Y160E}$ is 30,000-fold less than that for 20 nM PTP1B$^{1\text{-}321}$. (**C**) Measured $K_M$ and $k_{cat}/K_M$ values for PTP1B$^{1\text{-}321}$ alone (purple) and with 50 µM Grb2$^{Y160E}$ (blue) against all phosphopeptide substrates tested. (**D**) Dephosphorylation rates of the small molecule substrate *p*-nitrophenyl phosphate (pNPP) by PTP1B$^{1\text{-}321}$ with or without 50 µM Grb2$^{WT}$ as a function of pNPP concentration. (**E**) Dephosphorylation rates of pNPP by PTP1B$^{1\text{-}401}$ with or without 50 µM Grb2$^{WT}$ as a function of pNPP concentration.

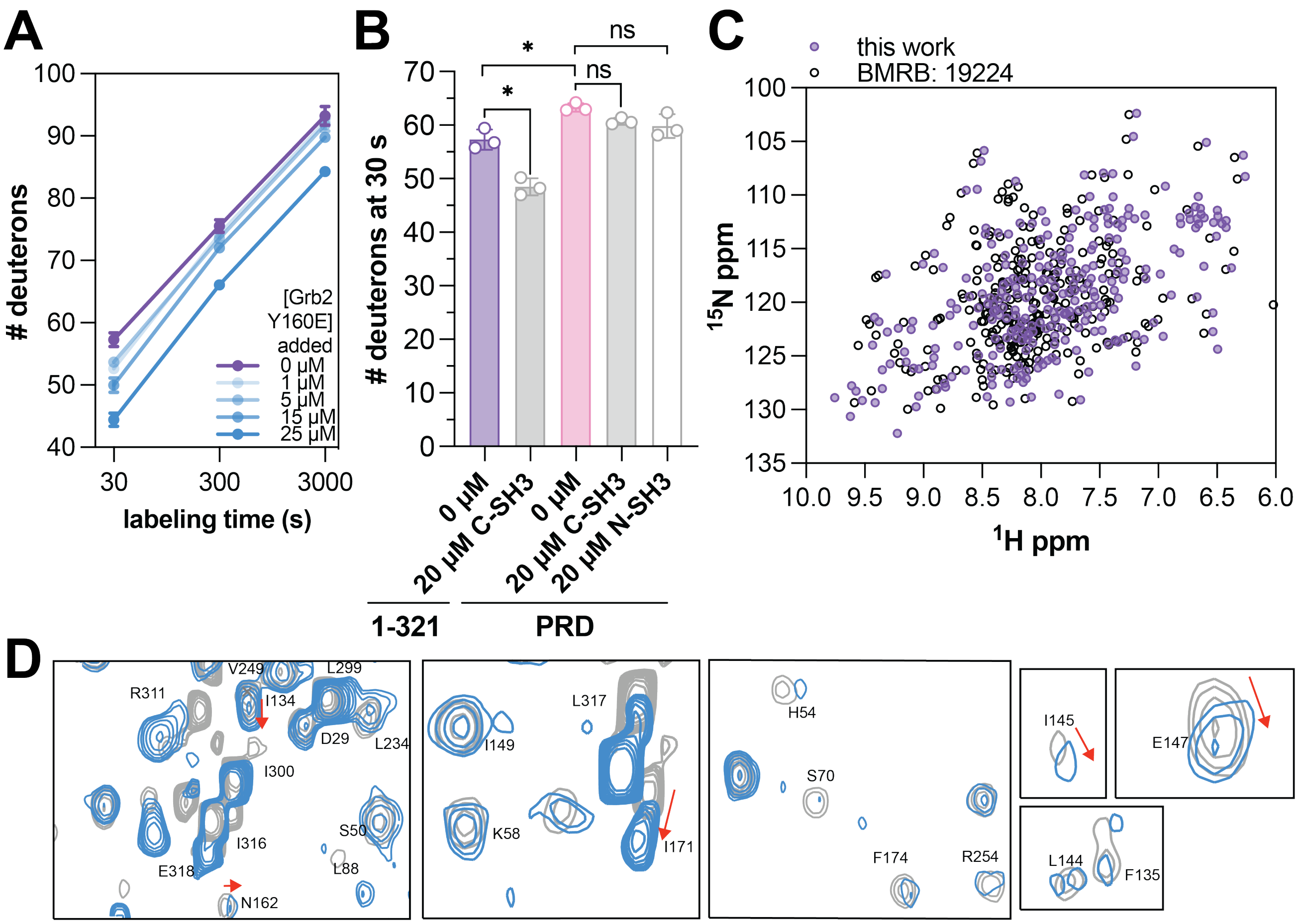


**Figure S4. Comparison of HDX-MS results and analysis of minor chemical shift changes observed by NMR.** (**A**) Deuterium exchange over time of PTP1B$^{1\text{-}321}$ in response to increasing Grb2$^{Y160E}$ concentrations. (**B**) Quantification of deuterium exchange of PTP1B$^{PRD}$ in response to isolated C- or N-SH3 domains. A paired, two-tailed t-test was used to test for significance (* denotes $p < 0.05$, ns denotes $p > 0.05$ or not significant). (**C**) Overlay of $^{1}$H-$^{15}$N HSQC spectra for PTP1B from this study (purple) with BMRB ID: 29224[4] (black circles). (**D**) Peaks corresponding to residues with minor changes, namely I134, I145, E147, N162, and I171, are indicated by red pointers. Peaks corresponding to residues H54, S70, I144, F174,R254 show considerable reduction in peak intensity after Grb2 binding.

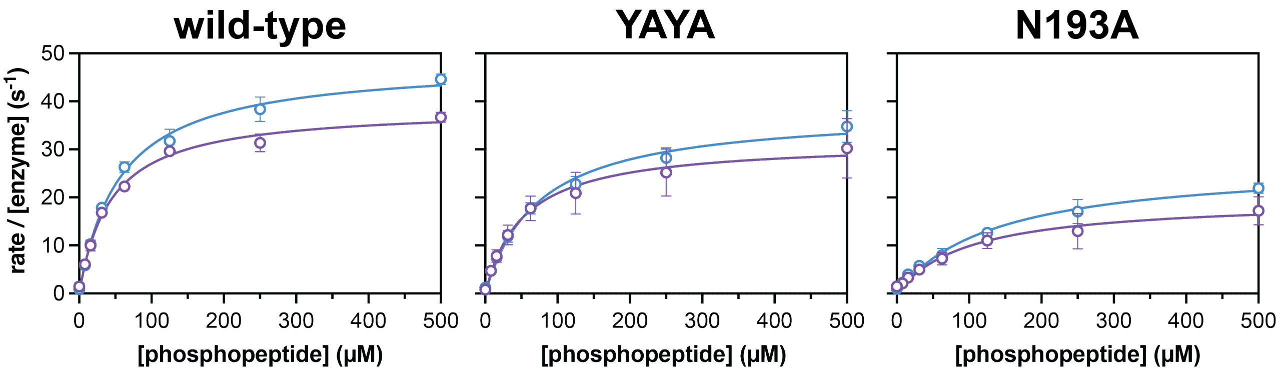


**Figure S5. Michaelis-Menten kinetics of PTP1B mutants with and without Grb2.** Kinetics of Gab1 pY627 dephosphorylation by wild-type PTP1B$^{1\text{-}321}$, the YAYA mutant, and the N193A mutant. Assays were carried out in the absence (purple) and presence Grb2$^{Y160E}$ (blue).

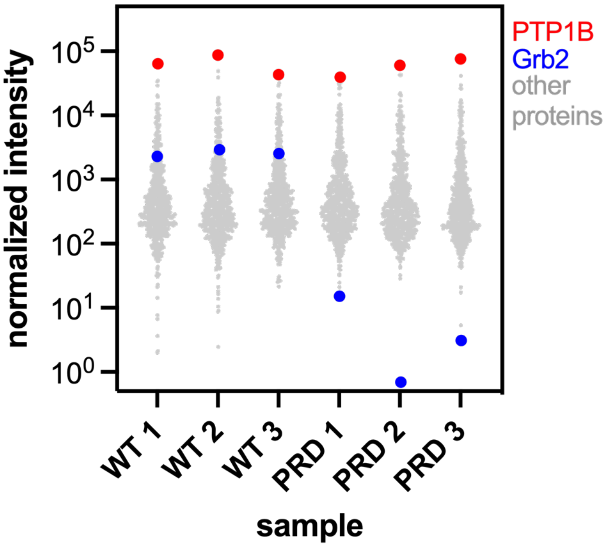


**Figure S6. Distribution of normalized intensities in MS proteomics replicates.** Raw intensity values for each protein group were adjusted by imputation and normalization, then filtered to remove proteins with fewer than five MS2 counts on average across either the $PTP1B^{WT}$ or $PTP1B^{PRD}$ replicates. The normalized intensities of the remaining proteins are plotted in gray, with PTP1B and Grb2 highlighted in red and blue, respectively.
